# Reproducible microbial community dynamics of two drinking water systems treating similar source waters

**DOI:** 10.1101/678920

**Authors:** Sarah C Potgieter, Zihan Dai, Minette Havenga, Solize Vosloo, Makhosazana Sigudu, Ameet J Pinto, Stefanus N Venter

## Abstract

Understanding whether the spatial-temporal dynamics of the drinking water microbiome are reproducible in full-scale drinking water systems is an important step towards devising engineering strategies to manipulate it. Yet, direct comparisons across full-scale drinking water systems are challenging because multiple factors, from source water to treatment process choice and configuration, can be unique to each system. This study compared the spatial-temporal dynamics of the drinking water microbiome in two drinking water treatment plants (DWTPs) with identical sequence of treatment strategies treating source waters from the same river system and with treated drinking water distributed in same large-scale (but independent) distribution system (DWDS) with similar disinfectant residual regiment. Dissimilarities in source water communities were tempered by the pre-disinfection treatments, resulting in highly similar post-filtration microbial communities between the two systems. However, high community turnover due to disinfection resulted in highly dissimilar microbial communities in the finished water between the two systems. Interestingly however, the microbial communities in the two systems increased in similarity during transit through the DWDS despite presence of a disinfectant residual. Overall our study finds that the drinking water microbiome demonstrated reproducible spatial and temporal dynamics within both independent but nearly identical DWTPs and their corresponding DWDSs.

**Graphical abstract:** 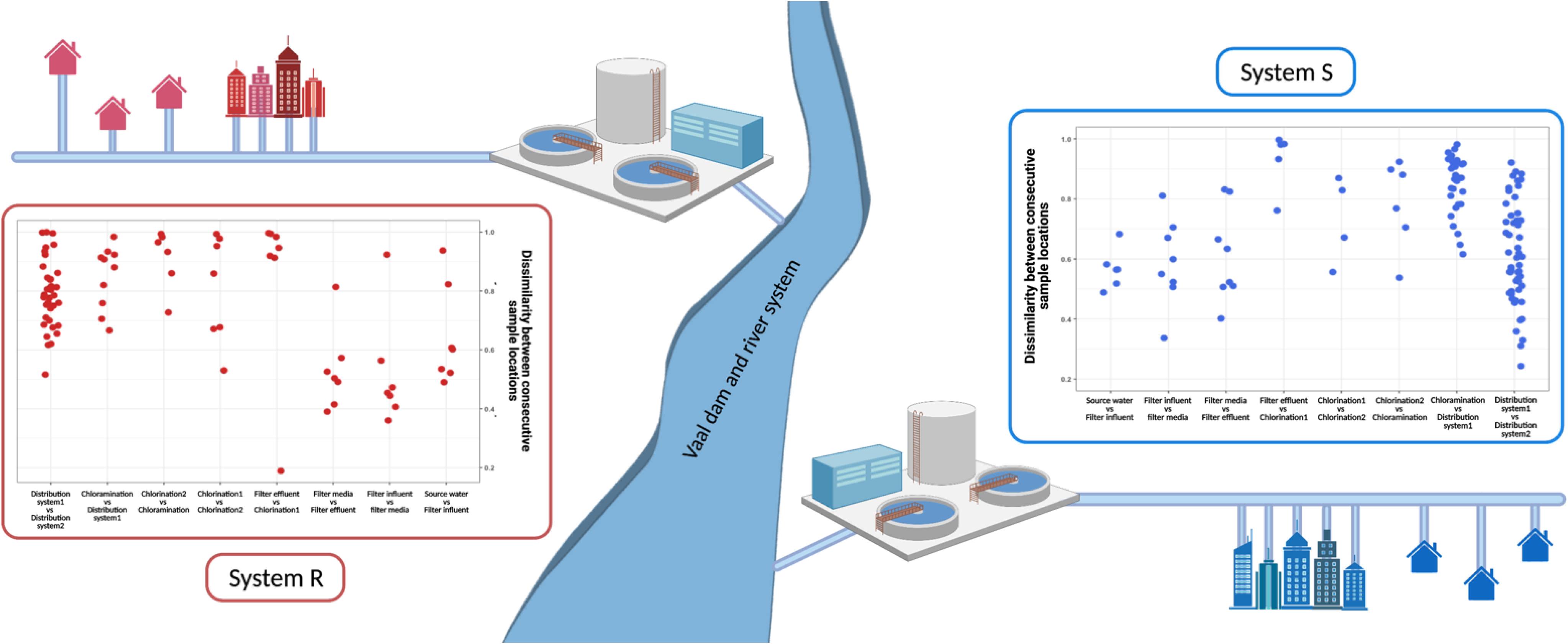

## 1. Introduction

Drinking water treatment operations are designed to reduce microbial concentrations and limit microbial growth in drinking water distribution systems (DWDS) in order to deliver safe water to the consumer^1–3^. The choice of treatment strategies implemented to achieve this is highly site specific and is based on a range of considerations including source water type, local regulations, etc^3,4^. Drinking water treatment itself can be considered as a series of ecological disturbances implemented sequentially on the microbial community as it transits from the source water to the consumers taps^5^. These disturbances shape the microbial community composition (i.e., who is there)^6^, as well as the population and community size (i.e., cell concentrations). Previous studies have shown that the drinking water microbiome is shaped by the choice of treatment strategy and conditions in the DWDS^3,5,7–10^. For instance, the microbial concentrations in finished drinking water can vary from 10^3^ and 10^5^ cells/mL^11–15^, depending on the presence, absence, and concentrations of the disinfectant residual. Disinfectant residuals not only impact community size, but also impact community structure^3,10,16–18^ and functional potential^18,19^. Similarly, filtration processes impact downstream microbial communities both by filtration-mediated seeding (Pinto *et al*., 2012) and through the biologically mediated removal of nutrients making them unavailable for microbial growth downstream^11,12,20,21^.

Factors influencing the drinking water microbiome are site specific due to the unique type and combination of treatment technologies across DWTPs and DWDS configurations, source water types, and operational practices^21–23^. Thus, while the impact of individual treatment process or environmental conditions on the drinking water microbiome may be compared across DWTP’s and DWDSs, it is often impossible to perform paired comparison between DWSs due to the inability to control for the type and sequential combination of treatment and distribution practices. For instance, studies have investigated drinking water microbiome dynamics in two DWTPs that use similar treatment processes but utilize different source waters or utilize the same source water but different treatment approaches^24–27^. Further, often such comparisons are limited to either the treatment system or the distribution system, but rarely encompass a comprehensive source-to-tap analysis. To our knowledge, no study has yet investigated the impact of identical drinking water treatment regimens and similar distribution practices on the drinking water microbiome in full-scale systems utilizing similar source waters. This is a significant knowledge gap in the field, because it has direct implications for our understanding of reproducibility of the drinking water microbiome dynamics and the impact of engineering interventions.

This study presents unique insights from systematic comparisons between two DWTPs, using the same treatment strategies to treat similar source waters originating from the same river system and their corresponding distribution systems. The specific objectives of this study were to investigate: (i) whether identical sequence of treatment technologies within two DWTP’s treating similar source waters results in similar changes in the microbial community, (ii) the extent to which similarities in temporal dynamics between DWTP’s are conserved (or not) between their respective distribution systems, and (iii) and to what extent the impact of physicochemical parameters on microbial community structure is conserved between the two DWSs. To our knowledge this mirrored study has not been previously performed and is critical towards improving our understanding of the spatial-temporal dynamics of the drinking water microbiome.

## 2. Materials and methods

### 2.1. Site description

This is study involves two drinking water systems (System R and S) that are operated by same drinking water utility (Fig. 1A). These two systems supply on an average 3653 million liters per day to approximately 12 million people within large metropolitan region and local municipalities as well as for industrial use through a network of large diameter pipelines stretching over 3056 km. The source water is drawn primarily from the Vaal river and dam system into two drinking water treatment plants (R_DWTP and S_DWTP), which abstract, treat, and distribute 98% (approximately 4320 ML/d) of the total water supplied by the utility. The R_DWTP (river intake pumping site) treats source water from the river downstream of the dam and the S_DWTP treats source water from a canal directly from the dam. Treatment of the source waters in both DWTPs consists of the identical treatment steps (Fig. 1B). Briefly, source water in both DWTPs is dosed with polyelectrolyte coagulants with low lime for coagulation and flocculation, with no need for pH correction after sedimentation. Although in some months in System R, a combination of polyelectrolyte and silica lime are used for coagulation and flocculation (Table S1). In these instances, following sedimentation, the water pH is adjusted to near neutral by bubbling CO2 gas followed by filtration through rapid gravity sand filters. Finally, the filter effluent is dosed with chlorine gas via bubbling as the primary disinfection step. The total chlorine at sites following chlorination varies between 1.0 mg/L and 2.5 mg/L after 20 min contact time. Chlorinated water leaving both DWTPs is again dosed with chloramine (approximately 2 mg/L) at secondary disinfection boosting stations. For the purpose of this study chlorinated water originating from the R_DWTP was monitored to the booster station, which supplies approximately 1,100 ML/d of chloraminated water, serving predominately the northwest area of the distribution system (R_DWDS). Chlorinated water originating from the S_DWTP was also monitored to another booster station, supplying approximately 700 ML/d of chloraminated water to the eastern parts of the distribution system (S_DWDS) (Fig. 1A). Samples were also collected after the booster stations in the chloraminated sections of each DWDS, with monochloramine residuals varying on average between 0.8 mg/L in the autumn and 1.5 mg/L in the spring. Further details on range of physical-chemical parameters for both systems were obtained from the utility (Table S1 and S2).

**Fig. 1:**
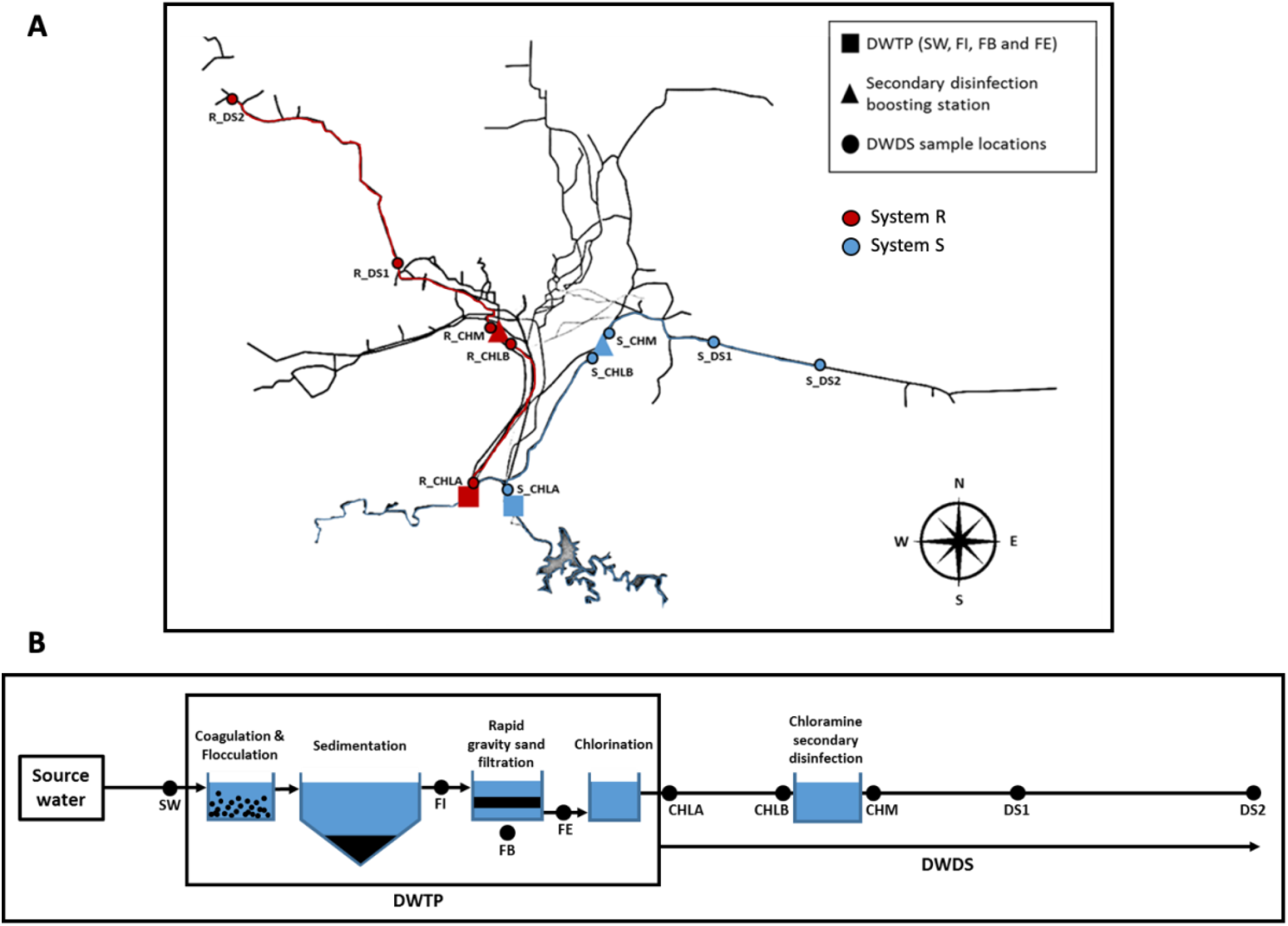
(A) Site map of the location of the drinking water treatment plants (R_DWTP and S_DWTP) and their corresponding distribution systems (R_DWDS and S_DWDS). System R is indicated in red and System S in blue. The two treatment plants are represented as squares, the two-secondary disinfection boosting stations, where chloramine is added, are represented as triangles and all sample locations are represented as circles. (B) Schematic of the layout of the DWTP and DWDS showing all sample locations sampled monthly for the duration of the study. Within the two DWTPs source water (SW), filter inflow (FI), filter bed media (FB) and filter effluent (FE) samples were collected. All other sample locations are indicated on the figure and described in the text.

### 2.2. Sample collection and processing

Samples were collected for 8 months (February 2016 – September 2016) on a monthly basis from the two DWTPs (R_DWTP and S_DWTP) and their associated DWDS (R_DWDS and S_DWDS) (Fig. 1A). At each DWTP, samples collected included source water (SW), filter influent (FI, i.e. water entering the rapid sand filter following coagulation, flocculation, sedimentation and carbonation), filter bed media (FB), filter effluent (FE) and chlorinated water leaving the treatment plant (CHLA). In both DWDSs, chlorinated water samples were collected immediately before the booster station (CHLB), chloraminated water leaving the booster station (CHM) and chloraminated bulk water samples at two points with the DWDSs (DS1 and DS2, respectively) (Fig. 1A and 1B). Within the two DWTPs, 1 L of source water, 4 L of filter influent, and 8 L of filter effluent were collected. Typically, 8 – 16 L of bulk water was collected for samples collected post disinfection and in the DWDS. Collected water samples were filtered to harvest microbial cells followed by phenol:chloroform DNA extraction as described by Potgieter *et al*., 2018. To obtain microbial biomass from the filter bed media samples, 10 g of filter media was mixed with 50 ml extraction buffer (i.e., 0.4 g/L EDTA, 1.2 g/L TRIS, 1 g/L peptone and 0.4 g/L N-dodecyl-N, N dimethyl-3-amminio-1-propanesulfonate) followed by sonication for 1 min to remove the microbial biomass attached to sand particles^28^. After sonication, the aqueous phase was filtered through a Sterivex™-GP 0.22 μm polycarbonate membrane filter unit (Merck Milipore, South Africa) followed by phenol:chloroform DNA extraction, as with bulk water samples.

### 2.3. Sequencing and data processing

Extracted DNA from samples were sent to the Department of Microbiology and Immunology, University of Michigan Medical School (Ann Arbor, USA) for the 2×250 bp sequencing of the V4 hypervariable region of 16S rRNA gene using the Illumina MiSeq platform^29^. All raw sequence data have been deposited with links to BioProject accession number PRJNA529765 on NCBI. The resultant 16S rRNA gene amplicon sequences were processed using the Divisive Amplicon Denoising Algorithm, DADA2 v1.14^30^ workflow including sequence filtering, dereplication, inferring sample composition, chimera identification and removal, merging of paired-end reads and construction on a sequence table. Initial trimming and filtering of reads followed standard filtering parameters described for Illumina MiSeq 2×250 V4 region of the 16S rRNA gene. Specifically, reads with ambiguous bases were removed (maxN=0) were removed, the maximum number of “expected errors” was defined (maxEE=2) and reads were truncated at the first instance of a quality score less than or equal to truncQ (truncQ=2). Dereplication was performed where identical sequences are combined into “unique sequences” while maintaining the corresponding abundance of the number of reads for that unique sequence. The core sample inference algorithm was applied to dereplicated data and forward and reverse reads were merged together to obtain fully denoised sequences^30^. Merged reads were then used to construct an amplicon sequence variant (ASV) table^31^, chimeras were identified and removed and taxonomic assignments were called using the SILVA reference database (https://www.arb-silva.de) through the DADA2 chimera removal and taxonomy assignment script.

### 2.4. Microbial community analysis

Resulting ASV table was imported into the mothur (v 1.35.1)^32^ and the shared sequences between sample locations from the two DWTPs and corresponding DWDSs as well as the unique sequences within each sample location were calculated using the venn function in mothur. Alpha diversity measures (i.e., richness, Shannon Diversity Index and Pielou’s evenness) were calculated using the summary.single function in mothur with the parameters, subsampling=1263 (sample with the least amount of sequences) and iters=1000 (1000 subsampling of the entire dataset). Due to subsampling, 10 samples were excluded from the analyses and Good’s coverage estimates were calculated to assess whether sufficient number of sequences were retained for each sample after subsampling. This indicated that subsampling at a library size of 1263 retained the majority of the richness for all samples (i.e., average Good’s coverage = 95.84 ± 0.02%). One-way analysis of variance (ANOVA)^33^ and post-hoc Tukey Honest Significant Differences (HSD) test were performed in R (http://www.R-project.org) using the stats package^34^ to determine the statistical significance between spatial and temporal groupings within the alpha diversity.

Bray-Curtis and weighted UniFrac were used to determine pair-wise dissimilarity in community structure between samples, whereas Jaccard and unweighted UniFrac were used to infer dissimilarity in community membership. Bray-Curtis and Jaccard distances were calculated using the dist.shared function in mothur with the parameters, subsampling=1263 and iters=1000. Weighted and unweighted UniFrac distances were calculated through the construction of a phylogenetic tree with representative sequences using the clearcut command in mothur also with the parameters subsampling=1263 and iters=1000^35,36^. Pairwise Analysis of Molecular Variance (AMOVA) was performed using the amova function in mothur on all beta diversity matrices, to determine the effect of sample groupings based on DWDS sample location, DWDS section, and season^37,38^. Beta diversity metrics and metadata files containing sample location, sample type, disinfection type and season were imported into R (http://www.R-project.org) for statistical analysis and visualization. Principal-coordinate analyses (PCoA) using Bray-Curtis and Jaccard distances was performed using the phyloseq package^39^. All plots were constructed using the ggplot2 package^40^. To identify the contribution of environmental parameters and their combinations towards microbial community structure, distance-based redundancy analysis (dbRDA) was used. The function dbrda() from R package “vegan”^41^ was applied on Bray-Curtis distances estimated between samples using ASV counts to investigate relationships between the scaled environmental parameters and microbial community structure. In addition, the faction of variation explained by the environmental parameters identified as significantly associated with ASV count-based Bray-Curtis distance matrices was determined using the function varpart() in the “vegan” package.

## 3. Results and discussion

A total of 172 samples were sequenced for this study (Table S3) which resulted in 4,921,399 sequences post quality control and a total of 10,012 ASVs. Taxonomic classification of these ASV’s revealed that bacteria dominated the microbial community (mean relative abundance, MRA: 98.74 ± 0.02% across all samples) followed by archaea (MRA: 1.04 ± 0.01%). The bacterial community was primarily composed of *Gammaproteobacteria* (including *Betaproteobacteriales*), *Actinobacteria*, and *Planctomycetes* (Fig. S1 and Table S4).

### 3.1. Treatment processes increased similarities between microbial communities across the two drinking water systems

Despite being drawn from the same surface water system, the two-source water microbial communities were significantly different (AMOVA, *F_ST_* ≤ 3.04, p < 0.001, depending on the beta diversity measure) and only shared 22.5% of the detected ASV’s (Fig. S2). These differences could likely be a result of varying hydrological conditions at the two locations (i.e., river drawn R_SW vs S_SW from the dam), which may translate to the variable occurrence of low abundance and transient taxa^42^. Despite harboring more ASV’s over the period of the study (Fig. S2), R_SW was less diverse than S_SW (ANOVA; p < 0.05) per timepoint (Fig. S3; Table S5). This was because R_SW was likely much more influenced by strong hydrological conditions such as runoff and increased flow rates during heavy rainfall events^43,44^ as compared to S_SW. This is supported by the fact that R_SW exhibited higher temporal variability as compared to S_SW based on pairwise beta diversity measures (Fig. S4).

The pre-chlorination treatment processes played an important role in tempering the differences in microbial communities across the two DWTPs. The microbial community composition of both DWTPs was highly diverse and in addition to *Proteobacteria* included other dominant phyla (i.e. MRA greater than 1%) such as *Acidobacteria, Cyanobacteria, Verrucomicrobia* and *Planctomycetes* (Fig. S1; Table S4). The 33 most abundant ASV’s i.e., ASV’s with an MRA of > 0.5%, showed similar trends between the two DWTPs in terms of increase/decrease in MRA with each sequential treatment step (Fig. 2; Table S6). These dominant ASV’s within both DWTP included members of *Actinobacteria, Acidobacteria, Gammaproteobacteria, Bacteroidetes* and *Thaumarcheota*. This reproducible effect of sequential treatment steps was also evident at the community level. Specifically, the treatment process resulted in similar changes in alpha diversity measures in both DWTPs (Fig. S3). Based on ANOVA these changes were found to be significant (richness: *F_ST_* = 19.67,*p* < 0.05, Shannon Diversity Index: *F_ST_* = 9.78, p < 0.05, Inverse Simpson Diversity Index: *F_ST_* = 15.64, p < 0.05 and Pielou’s evenness: *F_ST_* = 4.79, p < 0.05). Richness and diversity consistently decreased along treatment processes (excluding filter bed samples (FB)), with the most significant decrease immediately following chlorination (Fig. S3; Table S5). This shows that the decrease in microbial abundance and diversity, typical during treatment processes^1,5,45–47^, was reproducible in both systems.

**Fig. 2:**
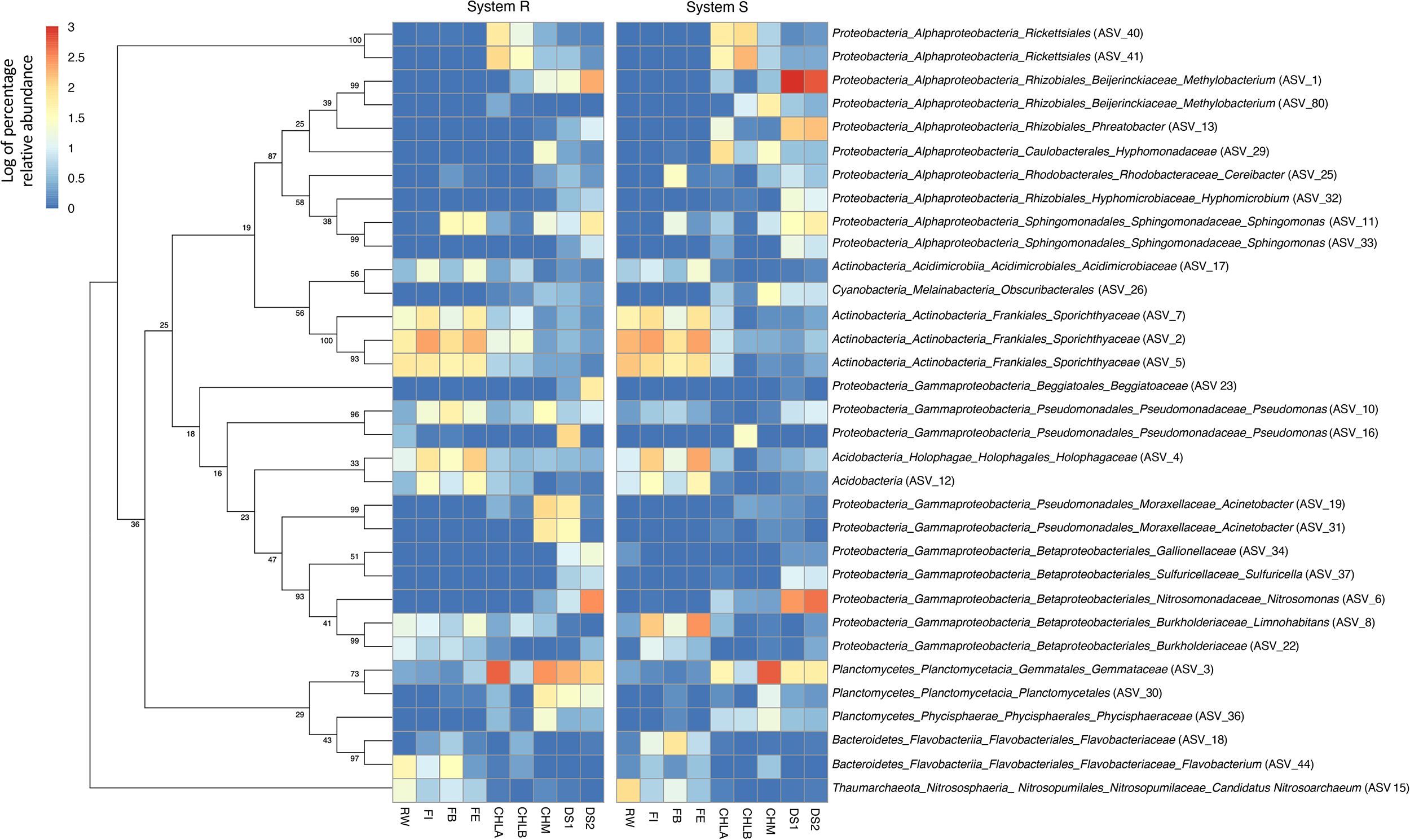
Maximum-likelihood phylogenetic tree (left) showing the groupings of the top ASV with a relative abundance > 0.5 (33 ASVs). Phylogenetic tree was constructed based on representative ASV DNA sequences with a bootstrap analysis of 1000 replicates (bootstrap values indicated as percentages). The log percentage of average relative abundance of those top ASVs for each sample site in each system are shown in heatmaps (right) followed by their taxonomic association. The average relative abundance for each sample location was averaged over duration of the study for each system. The log percentage relative abundance of each ASV is indicated in the legends on the left of the figure. See Table for mean relative abundances (MRA) of top dominant ASVs.

Beta-diversity analyses indicated that microbial communities became increasingly similar from source water through treatment and filtration where the microbial communities between the filter bed and the filter effluent were approximately 40% similar in community structure (Bray-Curtis: 0.57 ± 0.15) and 30% similar in community membership (Jaccard: 0.71 ± 0.11); this was comparable to the similarity of the filter bed and filter influent communities (community structure: Bray-Curtis: 0.55 ± 0.17, community membership: Jaccard: 0.7 ± 0.13) (Fig. 3A). Similar trends in community structure and membership were observed in weighted and unweighted Unifrac analyses (Fig. S5). Following similar prefiltration treatments of the source water, filter influent samples between the two systems showed the greatest similarity (Fig. 3B). Although, filter bed microbial communities (R_FB and S_FB) showed higher dissimilarity in community structure and membership (Fig. 3B; Table S7) between the two DWTPs, they were also significantly different from both the filter inflow (AMOVA: *F_ST_* ≤ 5.07, p < 0.001) and filter effluent (AMOVA: *F_ST_* ≤ 6.51, p < 0.001). The greater dissimilarity between the FB from the two DWTP’s could also be attributed to inherent high heterogeneity of attached growth microbial communities in the filter media^7,23^. Yet, microbial communities were more similar between filter bed samples of the two DWTPs, than the source waters that feed them. Further, an increase in the number of shared ASV’s between the two filter effluents (36.04%), indicates that conditions in the filter beds were sufficiently similar to have the same effect on the resulting effluent and the selection of dominant taxa in both systems.

**Fig. 3:**
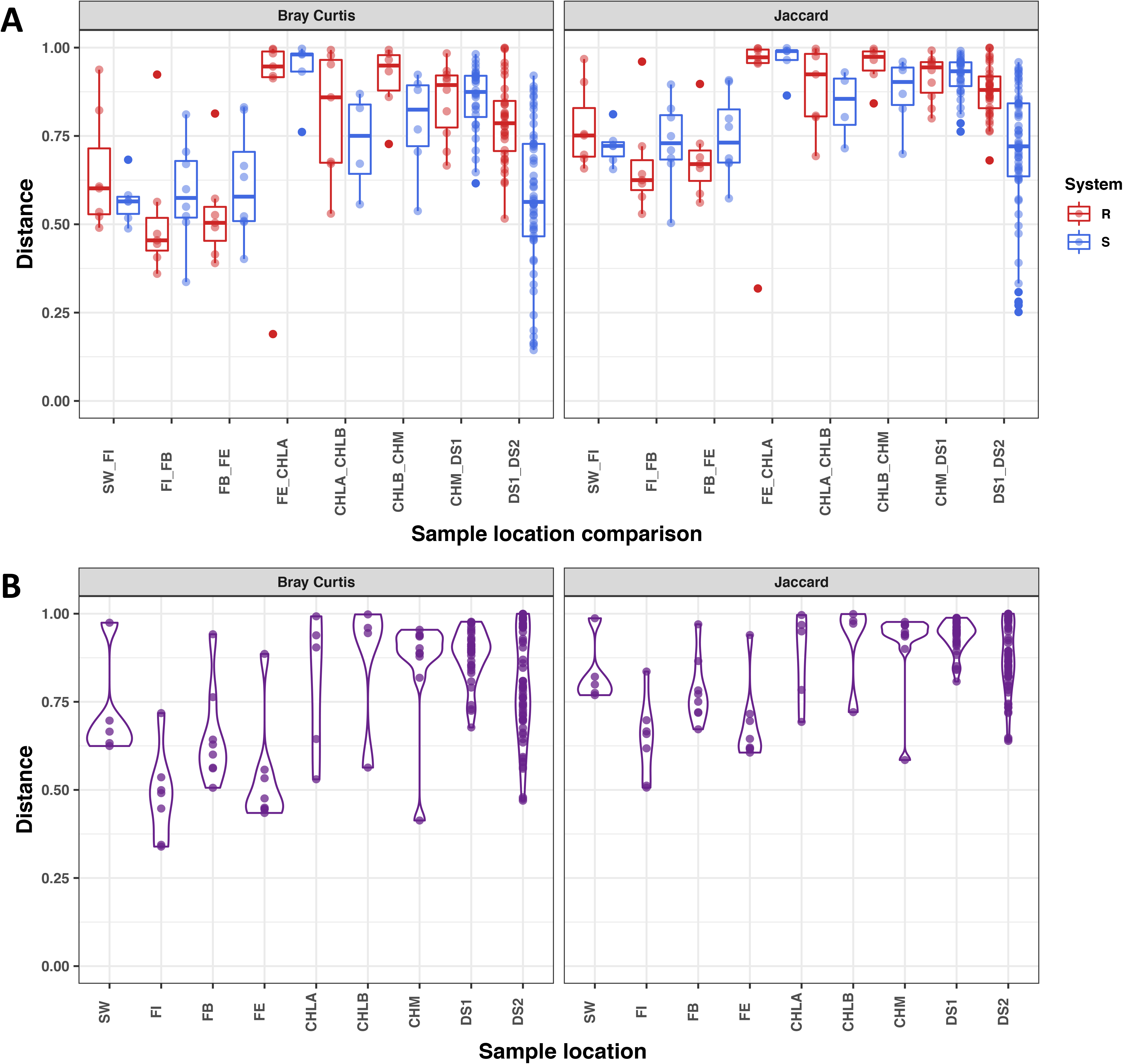
(A) Average pairwise beta diversity comparisons of both structure-based metric: Bray-Curtis and membership-based metric: Jaccard. Comparisons are between consecutive locations within each of the two systems for corresponding months. Sample abbreviations on the x-axis refer to comparisons of samples following the flow of bulk water through both systems, i.e. source water and filter inflow (SW_FI), filter inflow and filter bed media (FI_FB), filter bed media and filter effluent (FB_FE), filter effluent and chlorinated water leaving the DWTP (FE_CHLA), chlorinated water leaving the DWTP and chlorinated water entering the secondary disinfection boosting station (CHLA_CHLB), chlorinated water entering the secondary disinfection boosting station and chloraminated water (CHLB_CHM), chloraminated water and distribution system site 1 (CHM_DS1) and finally distribution system site 1 and distribution system site 2 (DS1_DS2). Sample comparisons from System R indicated in red and System S in blue. (B) Direct pairwise beta diversity comparisons (Bray-Curtis and Jaccard) between corresponding samples from the two systems. Pairwise beta diversity comparisons include samples from the same month. Sample abbreviations on the x-axis refer to source waters (SW), filter inflows (FI), filter bed medias (FB), filter effluents (FE), chlorinated waters leaving the DWTP (CHLA), chlorinated waters entering the secondary disinfection boosting station (CHLB), chloraminated waters (CHM), distribution system sites 1 (DS1) and distribution system sites 2 (DS2). Mean and standard deviations of each comparison is shown in Table S7. See Fig. S4 for corresponding weighted and unweighted Unifrac plots

### 3.2. Rapid microbial community turnover due to disinfection increases dissimilarity across the two drinking water systems

In contrast to the DWTP (i.e., pre-chlorination) where treatment processes enhance similarity in microbial community between the two DWTPs, the rapid change in microbial community due to chlorination resulted in significant increase in dissimilarity between locations within the DWTP/DWDS and across the two systems. Following chlorination, the microbial communities between filter effluent (FE) and bulk water (CHLA) were only 8 – 11% similar in community structure (Bray-Curtis: 0.89 ± 0.2) and membership (Jaccard: 0.92 ± 0.15) (AMOVA: *F_ST_* ≤ 18.22, p < 0.001 depending on the beta diversity measure) (Fig. 3A). This significant decrease in similarity was also observed in weighted and unweighted Unifrac analyses (Fig. S5). In addition, this significant change following disinfection was also observed as shift in the relative abundance of dominant ASVs (Fig. 2).

Further, chlorinated samples CHLA and CHLB increased slightly in similarity in community structure and membership (Bray-Curtis: 0.77 ± 0.16 and Jaccard: 0.86 ± 0.11) but remained highly dissimilar, though not significantly different. However, following secondary disinfection, the microbial communities between chlorinated (CHLB) and chloraminated water (CHM) again increased in dissimilarity in community structure (Bray-Curtis: 0.84 ± 0.12) and membership (Jaccard: 0.91 ± 0.08) (AMOVA: *F_ST_* ≤ 4.09, p < 0.001 depending on the beta diversity measure) in both systems (Fig. 3A) (same trends observed for Unifrac analyses, Fig. S5). This was consistent for both, the R and S systems. Beta diversity comparisons between paired samples from System R and S immediately after chlorination (R_CHLA and S_CHLA) showed an increase in dissimilarity in both community structure and membership (Fig. 3B; Table S7). This was also observed between R_CHLB and S_CHLB samples. Samples within the DWDS (CHM, DS1 and DS2) showed consistent temporal trends where samples from both systems increased in dissimilarity 6 months apart, consistent with the observed changes in temperature (Fig. S4; Fig. S6). However, temporal variability remained high within DS1 and DS2 samples where pair-wise comparisons between consecutive months within each sample location were dissimilar in community membership (i.e., Jaccard: 0.80 ± 0.06 and unweighted UniFrac: 0.67 ± 0.06) and structure (i.e., Bray-Curtis: 0.69 ± 0.13 and weighted UniFrac: 0.52 ± 0.13) (Fig. 3B).

Disinfection significantly reduces bacterial cell concentrations and has a substantial influence on community composition and structure ^3,10,24,45,48,49^. While microbial communities prior to chlorination were similar between the two DWTPs, the microbial community composition and structure in both systems were highly dissimilar post-chlorination. This dissimilarity was also observed on a temporal scale, where samples post disinfection showed greater temporal variability as compared to the pre-chlorinated samples. This variability could potentially arise from a few different factors. For instance, chlorine and chloramine are both strong oxidants and are likely to inactivate microorganisms indiscriminately; this could be one potential reason for higher temporal variability post-disinfection and greater dissimilarity between paired samples between the two drinking water systems. Alternatively, those surviving disinfection may be associated with biofilms. The presence of disinfectant residual has been reported to have limited impact in preventing biofilm development and in some cases may even promote biofilm formation as a stress response^50,51^. Another explanation could be reduction in microbial abundance and diversity due to disinfection may lead to the detection of low abundance or rare taxa that may not have well defined niche within the drinking water community and thus are variably present and/or detected.

Ultimately, the application of disinfection can be viewed as an ecological disturbance, where the sequential and controlled addition of disinfectants, disturb the drinking water microbial continuum^5^. The diversity of microbial communities generally decreases in response to environmental stress and disturbances, ultimately shifting the ecological balance of microbial populations within the community ^6,52,53^. Following a disturbance, the surviving populations are considered to have specific properties (such as biofilm formation or oxidative stress response mechanisms), allowing them to persist in the disrupted environment.

### 3.3. Chloramine residuals and distribution conditions promote microbial community stability

Considering the aforementioned similar impacts of treatment process and distribution system on the microbial communities between the two systems, it is not surprising that microbial communities from both systems clustered together depending on sampling location (i.e., DWTP or DWDS), rather than system (R or S) (Fig. 4). For instance, the DWTP locations clustered independently from the DWDS across both systems (Fig. 4A). Even within the DWTP, microbial communities clustered depending on sampling location despite the temporal variability (Fig. 4B). The same was consistent for the DWDS, where the chlorinated and chloraminated sections of the DWDS clustered together (Fig. 4C). Further, the measured water quality parameters were highly similar at each location between the two DWTPs and DWDS (Fig. S7; Table S2). To assess the extent to which measured water quality parameters impacted the microbial community, we performed dbRDA analysis by focusing primarily on the post-chlorination samples (i.e., chlorinated and chloraminated DWDS). This was done because each section of the DWDS across the two DWS’s has multiple samples per month over the eight-month sampling period resulting in significant sample size for both dbRDA and subsequent variance partitioning tests. Consistent with PCoA analyses, dbRDA revealed that samples clustered based on disinfectant residual type (chlorinated or chloraminated) and not based on the system they originated from. Based on dbRDA analysis, ammonium, water temperature, monochloramine and free Cl_2_ emerged as the best indicators among water quality parameters that explained the variability in microbial community structure in both systems (ANOVA: p < 0.001) (Fig. 5; Table S8). Increased concentrations of free chlorine (1.66 ± 0.55 mg/L) was identified as a significant variable in chlorinated samples (p = 0.0021).

**Fig. 4:**
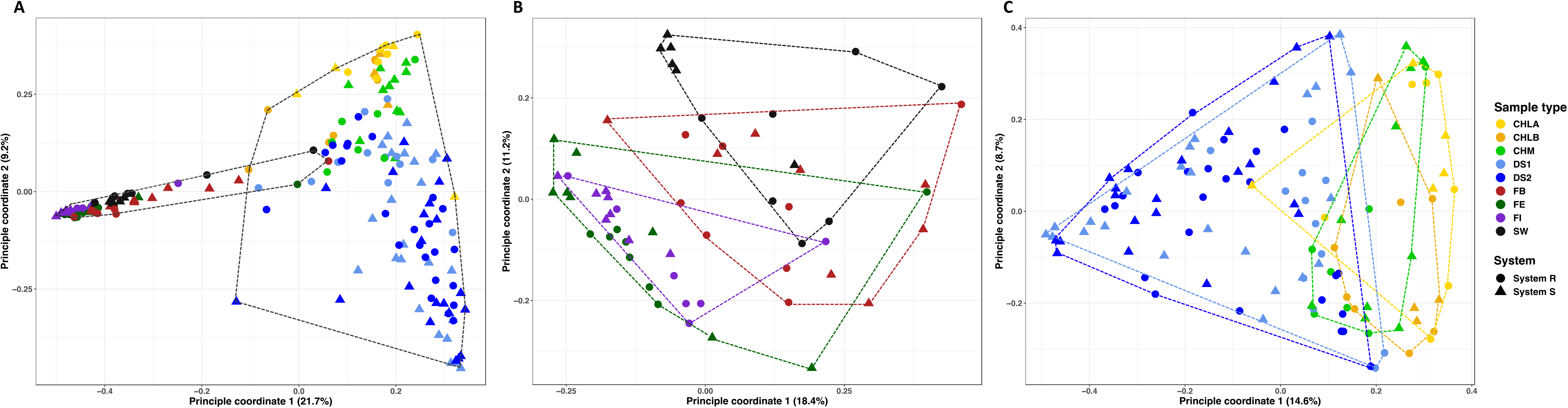
Principal coordinate analysis plot (based on Bray-Curtis dissimilarity) showing the spatial and temporal variability of the bacterial community structure among all samples from both systems (A), within the two DWTPs (B) and within the two corresponding DWDSs (C). Spatial groupings are shown where data points are colored based on sample location and shaped based on the system they originate from (System R samples are indicated as circles and System S samples as triangles). Color and shapes are indicated in the legends on the right of all plots.

**Fig. 5.**
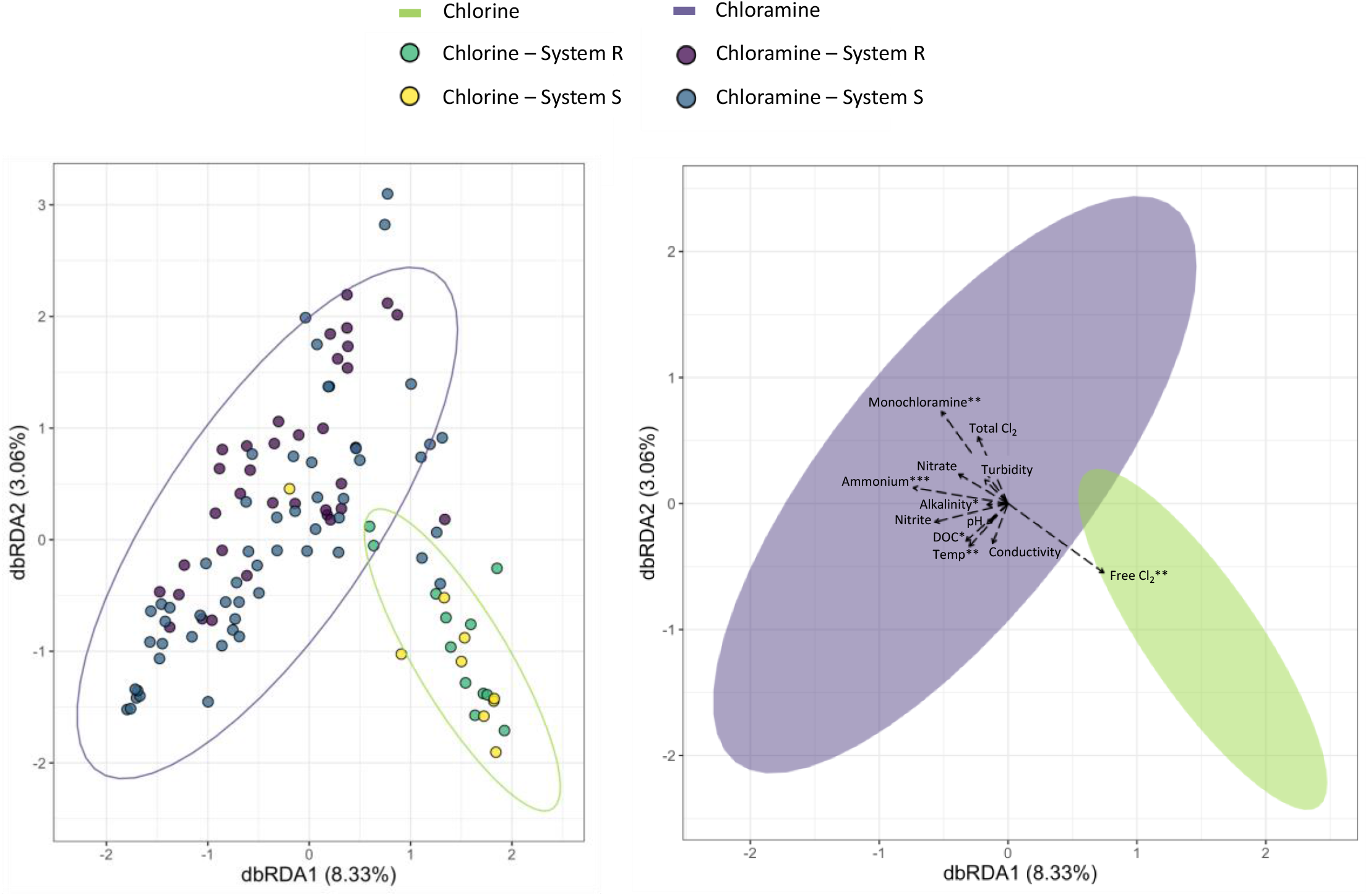
Relationship between measured water quality parameters and changes in the microbial community profile of locations distribution systems of the two systems where disinfectant residuals were applied. The plots represent the same distance-based redundancy analysis (dbRDA) based on Bray-Curtis distances of 16S rRNA amplicon sequencing data showing the clustering of individual samples (left) and their impacting water quality parameters (right). Chloraminated samples are were found to cluster in the purple ellipse and chlorinated samples in the green ellipse. Significance codes based on ANOVA are indicated as follows 0 ‘***’ 0.001 ‘**’ 0.01 ‘*’ 0.05 ‘.’ 0.1 ‘ ’ 1.

Alternatively, among other water quality parameters, ammonium (0.36 ± 0.13 mg/L as N) and monochloramine concentrations (1.60 ± 0.38 mg/L) as well as temperature (18.28 ± 4.47 °C) were identified as significant variables in chloraminated samples (p < 0.001). Variance partitioning analyses were used to determine the proportion of variance that could be attributed to variables (individual or combined) identified as most significant in the dbRDA analysis (Fig. 5). This resulted in ammonium, free Cl_2_, monochloramine and temperature each explaining 2.5%, 1.9%, 1.6% and 1.3% of the variance, respectively. While the contribution of these parameters towards explaining the variance in community structure was significant, it was small. In fact, 90% of the variance could not be explained by these four parameters (Fig. S8; Table S9). Here, ammonium and monochloramine concentrations and temperature best explained the variability in chloraminated samples from both distribution systems as disinfectant residual concentration and water temperatures did not differ greatly between the two lines. This is well supported as these variables are long considered to be important factors in shaping the drinking water microbiome^3^. Therefore, the observed dissimilarity between the two distribution lines may be accredited to the initial differential response of the microbial community to chlorine residuals.

Interestingly, the changes in the microbial community in the chlorinated sections were driven predominately by chlorine residuals (specifically free Cl_2_) as an initial significant ecological disturbance. However, chlorine residuals are typically short-lived and therefore chloramine residuals are added as a more stable alternative to ensure residuals are maintained over longer distances^54,55^. Nevertheless, the transport of bulk water over longer distances leads to increased water age and decreased disinfectant residual due to disinfectant decay, ultimately contributing to microbial regrowth^50^. In addition to disinfectant residuals, the microbial community is also exposed to various other distribution conditions and water quality parameters including (but not limited to) pipe material, hydraulic conditions, residence time, water temperature, concentrations of nitrogen species, organic carbon, etc. All these factors have a considerable impact on the drinking water microbiome^3^.

An increase in similarity was observed in locations towards the end of the DWDS (DS2 samples). Through distribution, water is continually seeded by similar microbial communities over time thereby selecting for the same dominant taxa through similarities in pipe material, residence times, hydraulic conditions and operation practices contributing to site specific taxa and biofilms. Biofilms have also been shown to seed bulk water communities through biofilm erosion, detachment and sloughing^56^. In this study, the established (historical) biofilms in both systems have been long exposed to similar distribution conditions thereby potentially contributing to the development of similar biofilm microbial communities. This increase in similarity in community membership and structure with increasing residence time in the DWDS was more pronounced in samples from summer and autumn. Here, elevated water temperatures in summer months may affect the bacterial community composition and structure by positively influencing the growth kinetics and competition processes of specific bacterial species in each section of the DWDS^3^. Alternatively, seasonal increase in water demand could increase shear stress resulting in increased detachment of similar established biofilms^56^.

In addition, where chlorine may be non-specific in its action in reducing bacterial cell concentrations, chloramine (with the addition of ammonia) may be more selective towards microbial taxa capable of using ammonia as an energy source. This may explain the observed increase in *Nitrosomonas* sp. (ASV_6) in chloraminated distribution system samples from both systems. Here, the addition of chloramine as a secondary disinfectant has been shown to support the growth of nitrifying bacteria in DWDS^57^. The long residence time and associated lower disinfectant residual concentrations, together with the release of ammonia through disinfection decay results in increased abundance of nitrifiers and therefore potential nitrification^49^. Thus, decreasing monochloramine residual concentrations associated with increased water age at nearing the end of both DWDSs may also contribute to the growth of similar bacterial assemblages at these sites.

This study provided a unique opportunity to compare the effects of the same treatment strategies (disturbances) on similar source waters as well as the distribution of treated water on the drinking water microbiome in a large-scale system. The spatial-temporal dynamics of two independent DWTP and their corresponding DWDSs with in the same large-scale distribution system were shown to be reproducible. Similarities in design and operational parameters of the two DWTPs resulted in the development of similar pre-disinfection microbial communities. However, the differential impact of chlorination was highlighted in post-disinfection samples from the two systems, resulting in dissimilar microbial communities between the two systems. Lastly, distribution was observed to select for certain dominant taxa, thereby increasing similarity between microbial communities due to the impact of inherent biofilms seeding the distribution system. However, dissimilarities in microbial community throughout distribution may arise from initial differences in the source waters and the differential response to chlorination ultimately leading to the presence of site specific rare/low abundant taxa. This study confirms the role of treatment and distribution in shaping the microbial community and may provide utilities with the reassurance that under that same conditions, drinking water quality will be reproducible throughout the varying stages of the drinking water system.

## Supporting Information

**Table S1:** Water quality parameters of the source water from both systems

**Table S2:** Water quality parameters for DWDS sites

**Table S3:** The number of samples collected and processed

**Table S4:** Mean relative abundance of dominant bacterial phyla

**Table S5:** Mean and standard deviations of alpha diversity indexes

**Table S6:** Mean percentage relative abundance of the top 33 abundant sequence variants

**Table S7:** Pair-wise beta diversity comparisons

**Table S8:** Permutation test for distance-based redundancy analysis (dbRDA)

**Table S9:** Variance Partition analyses

**Fig. S1:** Phylum-level and proteobacterial classes of bacterial sequences detected

**Fig. S2:** Venn diagram of shared amplicon sequence variants (ASVs) between source waters.

**Fig. S3:** Spatial changes in alpha diversity indexes

**Fig S4:** Temporal variation in beta diversity measures

**Fig S5:** Pairwise beta diversity comparisons of weighted and unweighted UniFrac analyses

**Fig S6:** Variation in temperature over the course of the study (8 months).

**Fig S7:** Variation in water quality parameters

**Fig S8:** Venn diagram based on Varpart analysis

## Acknowledgements

This research was funded and supported by Rand Water, Gauteng, South Africa through the Rand water Chair in Water Microbiology at the University of Pretoria. Sarah Potgieter would also like to acknowledge the National Research Foundation (NRF) for additional funding. Ameet J. Pinto was supported by NSF Award 1749530. Furthermore, the authors would like to acknowledge the Centre for Microbial Systems Molecular Biology Lab, University of Michigan, USA for their services in Illumina MiSeq sequencing.

## References

1. Hammes, F., Berney, M., Wang, Y., Vital, M., Koster, O. and Egli, T. (2008). Flow-cytometric total bacterial cell counts as a descriptive microbiological parameter for drinking water treatment processes. Water Research. 44(17): 4868–4877.

2. Lautenschlager, K., Hwang, C., Ling, F., Lui, W. T., Boon, N., Köster, O., Egli, T. and Hammes, F. (2014). Abundance and composition of indigenous bacterial communities in a multi-step biofiltration-based drinking water treatment. Water Research. 62: 40–52.

3. Prest, E. I., Hammes, F., van Loosdrecht, M. C. M. and Vrouwenvelder, J. S. (2016). Biological stability of drinking water: controlling factors, methods and challenges. Frontiers in Microbiology. 7(45): doi: 10.3389/fmicb.2016.00045.

4. Ho, L., Braun, K., Fabris, R., Hoefel, D., Morran, J., Monis, P. and Drikas, M., 2012. Comparison of drinking water treatment process streams for optimal bacteriological water quality. Water research, 46(12), pp.3934–3942.

5. Zhang, Y., Oh, S. and Liu, W. T. (2017). Impact of drinking water treatment and distribution on the microbiome continuum: an ecological disturbance's perspective. Environmental Microbiology. 19(8): 3163–3174.

6. Gomez-Alvarez, V., Pfaller, S., Pressman, J.G., Wahman, D.G. and Revetta, R.P., (2016). Resilience of microbial communities in a simulated drinking water distribution system subjected to disturbances: role of conditionally rare taxa and potential implications for antibiotic-resistant bacteria. Environmental Science: Water Research and Technology. 2(4): 645–657.

7. Pinto, A. J., Xi, C. and Raskin, L. (2012). Bacterial community structure in the drinking water microbiome is governed by filtration processes. Environmental Science and Technology. 46: 8851–8859.

8. Pinto, A., Schroeder, J., Lunn, M., Sloan, W. and Raskin, L. (2014). Spatial-temporal survey and occupancy-abundance modelling to predict bacterial community dynamics in the drinking water microbiome. mBIO. 5(3): e01135–14.

9. Bautista-de los Santos, Q. M., Schroeder, M. C., Sevillano-Rivera, M. C., Sungthong, R., Ijaz, U. Z., Sloan, W. T. and Pinto, A. J. (2016). Emerging investigators series: microbial communities in full-scale drinking water distribution systems – a meta-analysis. Environmental Science: Water Research and Technology. doi: 10.1039/c6ew00030d.

10. Potgieter, S., Pinto, A., Sigudu, M., Du Preez, H., Ncube, E. and Venter, S. (2018). Long-term spatial and temporal microbial community dynamics in a large-scale drinking water distribution system with multiple disinfectant regimes. Water Research. 139: 406–419.

11. Hammes, F., Berger, C., Koster, O. and Egli, T. (2010). Assessing biological stability of drinking water without disinfectant residuals in a full-scale water supply system. Journal of Water supply: Research and Technology-AQUA. 59: 31–40.

12. Lautenschlager, K., Hwang, C., Ling, F., Liu, W. T., Boon, N., Köster, O., Vrouwenvelder, H., Egli, T. and Hammes, F. (2013). A microbiology-based multi-parametric approach towards assessing biological stability in drinking water distribution networks. Water Research. 47: 3015–3025.

13. Liu, G., Verbeck, J. Q. J. C. and Van Dijk, J. C. (2013). Bacteriology of drinking water distribution systems: an integral and multidimensional review. Applied and Environmental Microbiology. 97: 9265–9276.

14. Gillespie, S., Lipphaus, P., Green, J., Parsons, S., Weir, P., Juskowiak, K., Jefferson, B., Jarvis, P. and Nocker, A. (2014). Assessing microbiological water quality in drinking water distribution systems with disinfectant residual using flow cytometry. Water Research. 65: 224–234.

15. Nescerecka, A., Rubulis, J., Vital, M., Juhna, T and Hammes, F. (2014). Biological instability in a chlorinated drinking water distribution network. PloS One. 9: e96354.

16. Gomez-Alvarez, V., Revetta, R. P. and Santo Domingo, J. W. (2012). Metagenomic analysis of drinking water receiving different disinfection treatments. Appied and Enivronmental Microbiology. 78(17): 6095–6102.

17. Hwang, C., Ling, F., Andersen, G. L., LeChevallier, M. W. and Liu, W. (2012). Microbial community dynamics of an urban drinking water distribution system subjected to phases of chloramination and chlorination treatments. Applied and Environmental Microbiology. 78(22): 7856–7865.

18. Dai, Z., Sevillano-Rivera, M.C., Calus, S.T., Bautista-de los Santos, Q.M., Eren, A.M., Van Der Wielen, P.W., Ijaz, U.Z. and Pinto, A.J., 2020. Disinfection exhibits systematic impacts on the drinking water microbiome. Microbiome, 8(1), pp.1–19.

19. Sevillano-Rivera, M., Dai, Z., Calus, S., Bautista-de Los Santos, Q.M., Eren, A.M., van der Wielen, P.W., Ijaz, U.Z. and Pinto, A.J., 2020. Differential prevalence and host-association of antimicrobial resistance traits in disinfected and non-disinfected drinking water systems. Science of the Total Environment, 749, p.141451.

20. Kirisits, M.J., Emelko, M.B. and Pinto, A.J., 2019. Applying biotechnology for drinking water biofiltration: advancing science and practice. Current opinion in biotechnology, 57, pp.197–204.

21. Ma, B., LaPara, T.M., N. Evans, A. and Hozalski, R.M., 2020. Effects of geographic location and water quality on bacterial communities in full-scale biofilters across North America. FEMS microbiology ecology, 96(2), p. fiz210.

22. Roeselers, G., Coolen, J., van der Wielen, P. W. J. J., Jaspers, M. C., Atsma, A., deGraaf, B. and Schuren, F. (2015). Microbial biogeography of drinking water: patterns in phylogenetic diversity across space and time. Environmental Microbiology. 17(7): 2505–2514.

23. Gülay, A., Musovic, S., Albrechtsen, H. J., Al-Soud, W. A., Sørensen, S. J. and Smets, B. F. (2016). Ecological patterns, diversity and core taxa of microbial communities in groundwater-fed rapid gravity filters. The ISME journal. 10(9): 2209.

24. Poitelon, J. B., Joyeux, M., Welté, B., Duguet, J. P., Prestel, E. and DuBow, M. S. (2010). Variations of bacterial 16S rDNA phylotypes prior to and after chlorination for drinking water production from two surface water treatment plants. Journal of Industrial Microbiology and Biotechnology. 37(2): 117–128.

25. Xu, J., Tang, W., Ma, J. and Wang, H., 2017. Comparison of microbial community shifts in two parallel multi-step drinking water treatment processes. Applied Microbiology and Biotechnology, 101(13), pp.5531–5541.

26. Nescerecka, A., Juhna, T. and Hammes, F. (2018). Identifying the underlying causes of biological instability in a full-scale drinking water supply system. Water Research. 135: 11–21.

27. Waak, M.B., Hozalski, R.M., Hallé, C. and LaPara, T.M., 2019. Comparison of the microbiomes of two drinking water distribution systems–with and without residual chloramine disinfection. Microbiome, 7(1), pp.1–14.

28. Camper, A. K., Lechevallier, M. W., Broadaway, S. C. and McFeters, G. A. (1985). Evaluation of procedures to desorb bacteria from granular activated carbon. Journal of Microbiological Method. 3: 187–198.

29. Kozich, J. J., Westcott, S. L., Baxter, N. T., Highlander, S. K., Schloss, P. D. (2013). Development of a dual-index strategy and curation pipeline for analyzing amplicon-sequencing data on the MiSeq Illumina sequencing platform. Applied and Environmental Microbiology. 79: 5112e5120.

30. Callahan, B. J., McMurdie, P. J., Rosen, M. J., Han, A. W., Johnson, A. J. A. and Holmes, S. P. (2016). DADA2: high-resolution sample inference from Illumina amplicon data. Nature Methods. 13(7): 581.

31. Callahan, B. J., McMurdie, P. J. and Holmes, S. P. (2017). Exact sequence variants should replace operational taxonomic units in marker-gene data analysis. The ISME journal. 11(12): 2639.

32. Schloss, P. D., Westcott, S. L., Ryabin, T., Hall, J. R., Hartmann, M., Hollister, E. B., Lesniewski, R. A., Oakley, B. B., Parks, D. H., Robinson, C. J., Sahl, J. W. (2009). Introducing mothur: open-source, platform-independent, community-supported software for describing and comparing microbial communities. Applied and Environmental Microbiology. 75(23): 7537e7541.

33. Chambers, J. M., Freeny, A. and Heiberger, R. M. (1992). Analysis of variance; designed experiments. Chapter 5 of statistical Models in S eds J. M. Chambers and T. J. Hastie, Wadsworth and Brooks/Cole.

34. R Core Team (2015). R: a language and environment for statistical computing. R Foundation for Statistical Computing, Vienna, Austria. http://www.R-project.org/.

35. Evans, J., Sheneman, L. and Foster, J.A. (2006). Relaxed neighbour-joining: a fast distance-based phylogenetic tree construction method. Journal of Molecular Evoltion. 62: 785e792.

36. Lozupone, C., Lladser, M. E., Knights, D., Stombaugh, J., Knight, R. (2011). UniFrac: an effective distance metric for microbial community comparison. The ISME Journal. 5(2):169e172.

37. Excoffier, L., (1993). Analysis of Molecular Variance (AMOVA) Version 1.55. Genetics and Biometry Laboratory, University of Geneva, Switzerland.

38. Anderson, M. J., (2001). A new method for non-parametric multivariate analysis of variance. Australian Ecology. 26 (1): 32e46.

39. McMurdie, P.J. and Holmes, S. (2013). Phyloseq: an R package for reproducible interactive analysis and graphics of microbiome census data. PLoS One. 8(4): e61217.

40. Wickham, H. (2009). ggplot2: Elegant graphics for data analysis. Springer-Verlag, New York. http://ggplot2.org.

41. Oksanen, J., Blanchet, F.G., Friendly, M., Kindt, R., Legendre, P., McGlinn, D., Minchin, P.R., O'Hara, R.B., Simpson, G.L., Solymos, P., Stevens, M.H.H., Szoecs, E., Wagner, H., 2019. Vegan: community ecology package.

42. Shade, A., Jones, S. E., Caporaso, J. G., Handelsman, J., Knight, R., Fierer, N., Gilbert, J. A. (2014). Conditionally rare taxa disproportionately contribute to temporal changes in microbial diversity. mBio ASM. 5(4): 1e9.

43. Prathumratana, L., Sthiannopkao, S. and Kim, K.W. (2008). The relationship of climatic and hydrological parameters to surface water quality in the lower Mekong River. Environment International. 34(6): 860–866.

44. Delpla, I., Jung, A.V., Baures, E., Clement, M. and Thomas, O. (2009). Impacts of climate change on surface water quality in relation to drinking water production. Environment International. 35(8): 1225–1233.

45. Lin, W., Yu, Z., Zhang, H. and Thompson, I. P. (2014). Diversity and dynamics of microbial communities at each step of treatment plant for potable water generation. Water Research. 52: 218–230.

46. Prest, E. I., El-Chakhtoura, J., Hammes, F., Saikaly, P.E., Van Loosdrecht, M.C.M. and Vrouwenvelder, J. S. (2014). Combining flow cytometry and 16S rRNA gene pyrosequencing: a promising approach for drinking water monitoring and characterization. Water Research. 63: 179–189.

47. Wang, H., Proctor, C. R., Edwards, M. A., Pryor, M., Santo Domingo, J. W., Ryu, H., et al. (2014). Microbial community response to chlorine conversion in a chloraminated drinking water distribution system. Environmental Science and Technology. 48: 10624–10633. doi: 10.1021/es502646d.

48. Eichler, S., Christen, R., Höltje, C., Westphal, P., Bötel, J., Brettar, I., Mehling, A. and Höfle, M. G. (2006). Composition and dynamics of bacterial communities of a drinking water supply system as assessed by RNA-and DNA-based 16S rRNA gene fingerprinting. Applied and Environmental Microbiology. 72(3): 1858–1872.

49. Wang, H., Masters, S., Edwards, M. A., Falkinham, J. O., and Pruden, A. (2014). Effect of disinfectant, water age, and pipe materials on bacterial and eukaryotic community structure in drinking water biofilm. Environmental Science and Technology. 48: 1426–1435. doi: 10.1021/es402636u

50. Fish, K. and Boxall, J. (2018). Biofilm Microbiome (Re) Growth Dynamics in Drinking Water Distribution Systems Are Impacted by Chlorine Concentration. Frontiers in Microbiology. 9: 2519.

51. Fish, K.E., Reeves-McLaren, N., Husband, S. and Boxall, J., 2020. Unchartered waters: the unintended impacts of residual chlorine on water quality and biofilms. npj Biofilms and Microbiomes, 6(1), pp.1–12.

52. Atlas, R.M., Horowitz, A., Krichevsky, M. and Bej, A.K., 1991. Response of microbial populations to environmental disturbance. Microbial Ecology, 22(1), pp.249–256.

53. Shade, A., Peter, H., Allison, S.D., Baho, D., Berga, M., Bürgmann, H., Huber, D.H., Langenheder, S., Lennon, J.T., Martiny, J.B. and Matulich, K.L., 2012. Fundamentals of microbial community resistance and resilience. Frontiers in microbiology, 3, p.417.

54. Neden, D.G., Jones, R.J., Smith, J.R., Kirmeyer, G.J. and Foust, G.W., 1992. Comparing chlorination and chloramination for controlling bacterial regrowth. Journal-American Water Works Association, 84(7), pp.80–88.

55. Norton, C. D and LeChevallier, M. W. (1997). Chloramination: its effect on distribution system water quality. J. AWWA. 89(7): 66–77.

56. Fish, K.E., Osborn, A.M. and Boxall, J., 2016. Characterising and understanding the impact of microbial biofilms and the extracellular polymeric substance (EPS) matrix in drinking water distribution systems. Environmental science: water research & technology, 2(4), pp.614–630.

57. Potgieter, S.C., Dai, Z., Venter, S.N., Sigudu, M. and Pinto, A.J., 2020. Microbial Nitrogen Metabolism in Chloraminated Drinking Water Reservoirs. mSphere, 5(2).

