## Supporting Information for "Reproducible microbial community dynamics of two drinking water systems treating similar source waters"

**APPENDIX A – Supplementary information**

**Supplementary tables**

**Table S1:** Water quality parameters of the source water from both systems

|  |  | **Source water quality** | | | **Treatment process** | |
| --- | --- | --- | --- | --- | --- | --- |
| **Sample** | **Date** | **Alkalinity** | **Turbidity (NTU)** | **pH** | **Lime Silica CO2 Ferric** | **Polymeric Coag lime (low)** |
| **System R** | | | | | | |
| R_SW1 | 2016/02/22 | 61 | 64 | 8.09 | 46% | 54% |
| R_SW2 | 2016/03/22 | 63 | 60 | 7.88 | 0% | 100% |
| R_SW3 | 2016/04/19 | 63 | 61 | 8.04 | 0% | 100% |
| R_SW4 | 2016/05/17 | 64 | 62 | 8.07 | 0% | 100% |
| R_SW5 | 2016/06/20 | 61 | 55 | 8.11 | 0% | 100% |
| R_SW6 | 2016/07/18 | 63 | 66 | 8.35 | 0% | 100% |
| R_SW7 | 2016/08/16 | 60 | 77 | 8.23 | 18% | 82% |
| R_SW8 | 2016/09/19 | 58 | 85 | 8.13 | 31% | 69% |
| **System S** | | | | | | |
| S_SW1 | 2016/02/01 | 58 | 58 | 8.2 | 0% | 100% |
| S_SW2 | 2016/03/07 | 67 | 57 | 8.3 | 0% | 100% |
| S_SW3 | 2016/04/04 | 59 | 62 | 8.3 | 0% | 100% |
| S_SW4 | 2016/05/09 | 58 | 67 | 8.5 | 0% | 100% |
| S_SW5 | 2016/06/06 | 59 | 58 | 8.7 | 0% | 100% |
| S_SW6 | 2016/07/04 | 57 | 71 | 8.5 | 0% | 100% |
| S_SW7 | 2016/08/01 | 57 | 75 | 8.8 | 0% | 100% |
| S_SW8 | 2016/08/29 | 51 | 71 | 8.7 | 0% | 100% |

**Table S2:** Water quality parameters for DWDS sites (CHLA – DS2) from both systems for the dates of sample collection (February – September 2016). Blank cells in the table indicate parameter data was not available at specific locations on the dates of sampling

| **System R** | | | | | | | | | | | | | |
| --- | --- | --- | --- | --- | --- | --- | --- | --- | --- | --- | --- | --- | --- |
| **Sample name** | **Month** | **Conductivity** | **DOC** | **Free Cl_2_** | **Total Cl_2_** | **NH_2_Cl** | **Alkalinity** | **NH_4_** | **NO_2_** | **NO_3_** | **pH** | **Temp** | **Turbidity** |
|  |  | **(mS/m)** | **(mg/l as C)** | **(mg/l)** | **(mg/l)** | **(mg/l)** | **(mg/l CaCO3)** | **(mg/l as N)** | **(mg/l as N)** | **(mg/l as N)** |  | **(°C)** | **(NTU)** |
| R_CHLA1 | FE | 20.1 | 3.09 | 2.2 |  |  | 65.6 | 0.05 |  | 0.08 | 8.08 | 24.7 | 0.272 |
| R_CHLA2 | Mar | 21.7 | 2.84 | 2.2 |  |  | 69.9 | 0.02 |  | 0.13 | 7.71 | 23.4 | 0.274 |
| R_CHLA3 | Apr | 20.7 | 3.4 | 2.08 |  |  | 70.9 | 0.02 |  | 0.11 | 8.17 | 18.3 | 0.273 |
| R_CHLA4 | May | 20.5 | 3.22 | 2.2 |  |  | 62.1 | 0.01 | 0 | 0.15 | 8.2 | 16.8 | 0.243 |
| R_CHLA5 | Jun | 20.9 | 2.63 | 2.06 |  |  | 69.8 | 0.01 |  | 0.18 | 8.02 | 13.2 | 0.26 |
| R_CHLA6 | Jul | 21.5 |  | 2.3 |  |  | 69 | 0.01 | 0 | 0.22 | 7.67 | 11.4 | 0.297 |
| R_CHLA7 | Aug | 20.7 |  | 2.12 |  |  | 68.2 |  | 0 | 0.24 | 8.22 | 13.4 | 0.292 |
| R_CHLA8 | Sept | 19 |  | 2.09 |  |  | 64.4 |  |  | 0.21 | 8.12 | 14.8 | 0.288 |
| R_CHLB1 | Feb | 26.2 |  | 1.09 | 1.13 | 0.04 | 81.7 |  |  |  | 7.64 | 24.1 | 0.305 |
| R_CHLB2 | Mar | 19.9 |  | 0.96 | 1.06 | 0.1 | 67.4 |  |  |  | 7.68 | 22.9 | 0.294 |
| R_CHLB3 | Apr | 21.3 |  | 1.15 | 1 | 0.05 | 69.5 |  |  |  | 8.07 | 20.9 | 0.281 |
| R_CHLB4 | May | 21.1 |  | 1.23 | 1.29 | 0.06 | 68.3 |  |  |  | 7.85 | 16.4 | 0.31 |
| R_CHLB5 | Jun | 21.1 |  | 1.18 | 1.22 | 0.04 | 75.2 |  |  |  | 8.31 | 15.2 | 0.264 |
| R_CHLB6 | Jul | 20.7 |  | 1.85 | 1.82 | 0.08 | 69.7 | 0.05 | 0 | 0.21 | 8.08 | 12.3 | 0.262 |
| R_CHLB7 | Aug | 19.5 |  | 1.31 | 1.36 | 0.05 | 60.3 |  | 0 | 0.24 | 8.15 | 13.6 | 0.338 |
| R_CHLB8 | Sept | 18.6 |  | 1.37 | 1.49 | 0.12 | 62.9 | 0.38 |  | 0.2 | 7.72 | 17.9 | 0.386 |
| R_CHM1 | Feb | 21.4 |  | 0.07 | 2.16 | 2.09 | 71.4 |  |  |  | 7.83 | 24.6 | 0.299 |
| R_CHM2 | Mar | 20.1 |  | 0.1 | 2.05 | 1.95 | 67 |  |  |  | 7.61 | 22.8 | 0.301 |
| R_CHM3 | Apr | 20.4 |  | 0.1 | 1.95 | 1.85 | 65.9 |  |  |  | 8.03 | 22.2 | 0.265 |
| R_CHM4 | May | 22.3 |  | 0.03 | 1.85 | 1.82 | 73.8 |  |  |  | 7.79 | 16.5 | 0.227 |
| R_CHM5 | Jun | 20.5 |  | 0.04 | 1.8 | 1.76 | 71.6 |  |  |  | 8.25 | 15 | 0.26 |
| R_CHM6 | Jul | 19.8 |  | 0.07 | 1.9 | 1.83 | 75.4 | 0.06 |  |  | 8.21 | 12.9 | 0.294 |
| R_CHM7 | Aug | 18.7 |  | 0.11 | 1.79 | 1.68 | 56.7 | 0.12 |  |  | 7.85 | 14.1 | 0.347 |
| R_CHM8 | Sept | 18.4 |  | 0.04 | 1.54 | 1.5 | 61.4 | 0.08 |  |  | 7.77 | 17.2 | 0.268 |
| R_DS1A1 | Feb | 21.9 | 2.81 | 0.05 | 1.6 | 1.55 | 71.8 | 0.27 | 0.06 | 0.09 | 7.98 | 24.4 | 0.221 |
| R_DS1A2 | Mar | 19.7 | 2.8 | 0.2 | 1.81 | 1.61 | 68.5 | 0.33 | 0.02 | 0.13 | 7.71 | 23.4 | 0.269 |
| R_DS1A3 | Apr | 25 | 3.223 | 0.18 | 1.84 | 1.67 | 76.4 | 0.3772 | 0.0051 | 0.07 | 7.79 | 21.2 | 0.26 |
| R_DS1A4 | May | 21.5 | 3.1912 | 0.05 | 1.92 | 1.87 | 66.6 | 0.4 | 0.0109 | 0.35 | 8.14 | 17.9 | 0.261 |
| R_DS1A5 | Jun | 21.5 | 2.974 | 0.26 | 2.2 | 1.94 | 74.4 | 0.39 | 0.0089 | 0.17 | 8.03 | 14.3 | 0.282 |
| R_DS1A6 | Jul | 17.7 |  | 0.06 | 1.91 | 1.85 | 66.2 | 0.36 | 0.01 | 0.3 | 8.17 | 13.5 | 0.254 |
| R_DS1A7 | Aug | 20.7 |  | 0.04 | 1.54 | 1.5 | 69.5 | 0.22 | 0 | 0.45 | 8.15 | 13.4 | 0.364 |
| R_DS1A8 | Sept | 18.8 |  | 0.2 | 1.84 | 1.64 | 63.7 | 0.35 |  | 0.24 | 8.4 | 16.1 | 0.306 |
| R_DS1B1 | Feb | 21.2 | 2.85 | 0.08 | 1.97 | 1.89 | 73.5 | 0.29 | 0.01 | 0.16 | 7.99 | 24.5 | 0.308 |
| R_DS1B2 | Mar | 20.3 | 2.71 | 0.58 | 1.32 | 0.74 | 101.6 | 0.37 | 0.01 | 0.2 | 7.72 | 23.8 | 0.359 |
| R_DS1B3 | Apr | 22 | 3.22 | 0.56 | 1.9 | 1.34 | 73.3 | 0.44 | 0.01 | 0.14 | 8.11 | 21.5 | 0.239 |
| R_DS1B4 | May | 21.4 | 3.27 | 0.08 | 1.95 | 1.87 | 66.4 | 0.4 | 0.01 | 0.08 | 8.02 | 17.8 | 0.19 |
| R_DS1B5 | Jun | 21.3 | 3.43 | 0.16 | 2.05 | 1.89 | 73.4 | 0.39 | 0.01 | 0.17 | 8.02 | 14.5 | 0.318 |
| R_DS1B6 | Jul | 17.5 |  | 0.07 | 1.75 | 1.68 | 67.4 | 0.35 | 0.01 | 0.29 | 8.1 | 14.5 | 0.347 |
| R_DS1B7 | Aug | 20.8 |  | 0.02 | 1.67 | 1.65 | 68.9 | 0.26 | 0 | 0.31 | 8.13 | 13.4 | 0.269 |
| R_DS1B8 | Sept | 18.8 |  | 0.17 | 1.84 | 1.67 | 66.8 | 0.32 |  | 0.25 | 8.19 | 16.1 | 0.32 |
| R_DS2A1 | Feb | 21.6 | 2.97 | 0.05 | 1.33 | 1.28 | 70.9 | 0.24 | 0.2 | 0.19 | 8.12 | 25.8 | 0.21 |
| R_DS2A2 | Mar | 20.6 | 3.62 | 0.72 | 1.44 | 0.72 | 68.8 | 0.24 | 0.15 | 0.18 | 7.86 | 23.6 | 0.328 |
| R_DS2A3 | Apr | 21.1 | 2.68 | 0.63 | 1.66 | 1.03 | 67.9 | 0.39 | 0.09 | 0.31 | 7.74 | 22.3 | 0.265 |
| R_DS2A4 | May | 21.7 | 3.29 | 0.05 | 1.69 | 1.64 | 72 | 0.67 | 0.02 | 0.7 | 8.18 | 19.4 | 0.264 |
| R_DS2A5 | Jun | 21 | 3.12 | 0.13 | 2.01 | 1.88 | 76.1 | 0.42 | 0.02 | 0.51 | 8.19 | 16.9 | 0.237 |
| R_DS2A6 | Jul | 19.5 |  | 0.41 | 1.85 | 1.44 | 69.9 | 0.35 | 0.01 | 0.26 | 8.24 | 14.3 | 0.349 |
| R_DS2A7 | Aug | 21.4 |  | 0.05 | 1.53 | 1.48 | 65 | 0.23 | 0 | 0.44 | 8.19 | 14.5 | 0.267 |
| R_DS2A8 | Sept | 19 |  | 0.11 | 1.48 | 1.37 | 60.7 | 0.31 |  | 0.17 | 7.97 | 19.1 | 0.313 |
| R_DS2B1 | Feb | 21.4 | 3.31 | 0.07 | 1.27 | 1.2 | 70.3 | 0.28 | 0.17 | 0.2 | 8.05 | 25.8 | 0.211 |
| R_DS2B2 | Mar | 20.5 | 2.71 | 0.65 | 1.21 | 0.56 | 68.1 | 0.21 | 0.16 | 0.18 | 7.72 | 23.5 | 0.291 |
| R_DS2B3 | Apr | 20.3 | 2.72 | 0.26 | 1.65 | 1.39 | 74.3 | 0.42 | 0.07 | 0.1 | 7.88 | 22.3 | 0.338 |
| R_DS2B4 | May | 21.7 | 3.3 | 0.02 | 1.62 | 1.6 | 70.6 | 0.76 | 0.02 | 0.19 | 8.04 | 19.7 | 0.227 |
| R_DS2B5 | Jun | 21.1 | 3.08 | 0.04 | 1.96 | 1.92 | 76.4 | 0.41 | 0.02 | 0.19 | 8.11 | 16.8 | 0.247 |
| R_DS2B6 | Jul | 19.4 |  | 0.14 | 1.67 | 1.53 | 68.9 | 0.36 | 0.01 | 0.29 | 8.13 | 14.4 | 0.373 |
| R_DS2B7 | Aug | 20.8 |  | 0.05 | 1.57 | 1.52 | 63.1 | 0.27 | 0 | 0.25 | 8.11 | 14.4 | 0.251 |
| R_DS2B8 | Sept | 18.8 |  | 0.1 | 1.57 | 1.47 | 69.6 | 0.31 |  | 0.19 | 7.79 | 18.7 | 0.29 |
| R_DS2C1 | Feb | 21.5 |  | 0.04 | 1.18 | 1.14 | 71.2 |  |  |  | 8.12 | 25.9 | 0.209 |
| R_DS2C2 | Mar | 21 |  | 0.26 | 1.3 | 1.04 | 67.6 | 0.26 | 0.17 | 0.35 | 7.85 | 23.9 | 0.344 |
| R_DS2C3 | Apr | 20.3 |  | 0.23 | 1.63 | 1.4 | 66.4 |  |  |  | 8.06 | 21.6 | 0.289 |
| R_DS2C4 | May | 21.5 |  | 0.12 | 1.75 | 1.63 | 72 |  |  |  | 8.16 | 18.6 | 0.316 |
| R_DS2C5 | Jun | 20.1 |  | 0.09 | 1.9 | 1.81 | 71.9 | 0.39 |  | 0.16 | 8.06 | 17.9 | 0.249 |
| R_DS2C6 | Jul | 18 |  | 0.02 | 1.83 | 1.81 | 66.3 | 0.37 | 0.17 | 0.21 | 8.12 | 13.8 | 0.266 |
| R_DS2C7 | Aug | 20.9 |  | 0.06 | 1.62 | 1.56 | 64 | 0.1 | 0.01 | 0.3 | 8.14 | 14.1 | 0.364 |
| R_DS2C8 | Sept | 19 |  | 0.12 | 1.51 | 1.39 | 59.9 | 0.31 | 0 | 0.2 | 7.93 | 18.1 | 0.257 |
| **System S** | | | | | | | | | | | | | |
| **Sample name** | **Month** | **Conductivity** | **DOC** | **Free Cl_2_** | **Total Cl_2_** | **NH_2_Cl** | **Alkalinity** | **NH_4_** | **NO_2_** | **NO_3_** | **pH** | **Temp** | **Turbidity** |
|  |  | **(mS/m)** | **(mg/l as C)** | **(mg/l)** | **(mg/l)** | **(mg/l)** | **(mg/l CaCO3)** | **(mg/l as N)** | **(mg/l as N)** | **(mg/l as N)** |  | **(°C)** | **(NTU)** |
| S_CHLA1 | Feb |  |  |  | 2.1 |  |  |  |  |  | 8.25 |  | 0.22 |
| S_CHLA2 | Mar |  |  |  | 2.1 |  |  |  |  |  | 8.25 |  | 0.18 |
| S_CHLA3 | Apr | 17.7 |  | 2 | 2.3 |  |  |  |  |  | 8.6 | 21.09 | 0.12 |
| S_CHLA4 | May | 19.3 |  | 1.85 | 1.94 |  |  |  |  |  | 8.3 | 18.83 | 0.13 |
| S_CHLA5 | Jun | 19.9 |  | 2 | 1.85 |  |  |  |  |  | 8.4 | 16.06 | 0.14 |
| S_CHLA6 | Jul | 19 |  | 2.25 | 1.99 |  |  |  |  |  | 7.99 | 14.58 | 0.26 |
| S_CHLA7 | Aug | 19.3 |  | 2.1 | 2.13 |  |  |  |  |  | 8.35 | 14.57 | 0.15 |
| S_CHLA8 | Sept |  |  |  | 1.9 |  |  |  |  |  |  |  |  |
| S_CHLB1 | Feb |  |  | 1.08 | 1.1 | 0.02 |  |  |  |  |  | 24.6 | 0.3 |
| S_CHLB2 | Mar |  |  | 1.2 | 1.25 | 0.05 |  |  |  |  |  | 24.2 | 0.25 |
| S_CHLB3 | Apr | 18.3 |  | 1.14 | 1.25 | 0.11 |  |  |  |  | 7.85 | 22 | 0.27 |
| S_CHLB4 | May |  |  | 1.16 | 1.22 | 0.06 |  |  |  |  |  | 18.4 | 0.35 |
| S_CHLB5 | Jun |  |  | 0.77 | 0.84 | 0.07 |  |  |  |  |  | 16 | 0.25 |
| S_CHLB6 | Jul | 18 |  | 1.2 | 1.18 | 0.13 |  | 0.11 | 0.03 | 0.15 | 8.24 | 11.7 | 0.15 |
| S_CHLB7 | Aug | 22 |  | 1.39 | 1.39 | 0.14 | 71 | 0.05 | 0.03 | 0.29 | 7.85 | 11.3 | 0.26 |
| S_CHLB8 | Sept | 17 |  | 1.3 | 1.27 | 0.04 |  | 0.05 | 0.03 | 0.38 | 7.98 | 14.1 | 0.24 |
| S_CHMA1 | Feb | 19 | 3.5 | 0.06 | 2.28 | 2.22 | 77 | 0.287 | 0.03 | 0.1 | 8.31 | 25.9 | 0.32 |
| S_CHMA2 | Mar | 19 | 2.8 | 0.07 | 2.44 | 2.37 | 70 | 0.243 | 0.03 | 0.1 | 7.85 | 25 | 0.25 |
| S_CHMA3 | Apr | 22 | 4 | 0.05 | 2.16 | 2.11 | 75 | 0.36 | 0.03 | 0.13 | 8.13 | 21.2 | 0.32 |
| S_CHMA4 | May | 19 | 3.3 | 0.06 | 2.34 | 2.28 | 80 | 0.47 | 0.03 | 0.1 | 8.12 | 19.1 | 0.31 |
| S_CHMA5 | Jun | 21 | 3.1 | 0.07 | 2.2 | 2.13 | 74 | 0.37 | 0.03 | 0.18 | 8.03 | 16.1 | 0.27 |
| S_CHMA6 | Jul | 18 |  | 0.01 | 2.2 | 2.19 |  |  | 0.03 | 0.23 | 8.24 | 12.3 | 0.2 |
| S_CHMA7 | Aug | 21 |  | 0.06 | 2.02 | 1.96 | 69 |  | 0.03 | 0.28 | 7.98 | 12.4 | 0.32 |
| S_CHMA8 | Sept | 17 |  | 0.07 | 2.16 | 2.09 |  |  | 0.03 | 0.22 | 8 | 15.7 | 0.22 |
| S_CHMB2 | Mar | 20 | 2.8 | 0.06 | 1.94 | 1.88 | 72 | 0.27 | 0.03 | 0.1 | 7.88 | 21.7 | 0.25 |
| S_CHMB3 | Apr | 23 | 3.3 | 0.05 | 2.18 | 2.13 | 79 | 0.3 | 0.03 | 0.15 | 7.91 | 21.4 | 0.31 |
| S_CHMB4 | May | 19 | 3 | 0.1 | 2.1 | 2 | 79 | 0.48 | 0.03 | 0.17 | 8.12 | 18.1 | 0.34 |
| S_CHMB5 | Jun | 21 | 3.1 | 0.05 | 2.12 | 2.07 | 74 | 0.37 | 0.03 | 0.22 | 8.42 | 16 | 0.25 |
| S_CHMB6 | Jul | 18 | 2.2 | 0.05 | 2.2 | 2.15 | 80 |  | 0.03 | 0.24 | 8.23 | 12 | 0.48 |
| S_CHMB7 | Aug | 21 | 2.1 | 0.05 | 2.08 | 2.03 | 69 |  | 0.03 | 0.28 | 7.9 | 11.4 | 0.28 |
| S_CHMB8 | Sept |  |  | 0.05 | 2.12 | 2.07 |  |  |  |  |  | 15.6 | 0.39 |
| S_DS1A1 | Feb | 27 | 3.3 | 0.05 | 1.2 | 1.15 | 85 | 0.19 | 0.11 | 0.2 | 7.62 | 25.2 | 0.4 |
| S_DS1A2 | Mar | 22 | 2.7 | 0.09 | 1.86 | 1.77 | 74 | 0.18 | 0.03 | 0.19 | 7.62 | 23.4 | 0.38 |
| S_DS1A3 | Apr | 20 | 3.7 | 0.01 | 1.54 | 1.53 | 72 | 0.37 | 0.03 | 0.24 | 7.96 | 21.6 | 0.29 |
| S_DS1A4 | May | 20 | 3.1 | 0.05 | 1.72 | 1.67 | 85 | 0.54 | 0.03 | 0.17 | 8.05 | 17 | 0.33 |
| S_DS1A5 | Jun | 19 | 2.6 | 0.05 | 1.79 | 1.74 | 77 | 0.49 | 0.03 | 0.17 | 8.34 | 15.9 | 0.24 |
| S_DS1A6 | Jul | 17 |  | 1.23 | 2.06 | 0.83 |  | 0.52 | 0.03 | 0.25 | 8 | 13.1 | 0.23 |
| S_DS1A7 | Aug | 17 |  | 0.18 | 2.03 | 1.85 |  | 0.47 | 0.03 | 0.27 | 8.18 | 12.3 | 0.32 |
| S_DS1A8 | Sept | 17 |  | 0.2 | 2.02 | 1.82 |  | 0.53 | 0.03 | 0.41 | 7.9 | 15 | 0.27 |
| S_DS1B1 | Feb | 19 | 3.3 | 0.06 | 1.13 | 1.07 | 72 | 0.2 | 0.11 | 0.23 | 8.09 | 25.6 | 0.25 |
| S_DS1B2 | Mar | 22 | 2.7 | 0.05 | 1.73 | 1.68 | 74 | 0.31 | 0.03 | 0.12 | 7.61 | 23.5 | 0.25 |
| S_DS1B3 | Apr | 21 | 3.6 | 0.02 | 1.76 | 1.74 | 76 | 0.39 | 0.03 | 0.13 | 7.98 | 21.5 | 0.28 |
| S_DS1B4 | May | 20 | 3.1 | 0.07 | 1.75 | 1.68 | 84 | 0.53 | 0.03 | 0.1 | 8.03 | 16.9 | 0.34 |
| S_DS1B5 | Jun | 20 | 2.6 | 0.04 | 1.74 | 1.7 | 78 | 0.45 | 0.03 | 0.2 | 8.19 | 15.2 | 0.31 |
| S_DS1B6 | Jul | 11 |  | 1.9 | 2 | 0.1 |  | 0.49 | 0.03 | 0.21 | 8.19 | 13.1 | 0.27 |
| S_DS1B7 | Aug | 17 |  | 0.33 | 2.01 | 1.68 |  | 0.45 | 0.03 | 0.28 | 8.15 | 12.6 | 0.44 |
| S_DS1B8 | Sept | 17 |  | 0.17 | 2.12 | 1.95 |  | 0.54 | 0.03 | 0.25 | 7.93 | 15 | 0.23 |
| S_DS1C1 | Feb | 20 | 3.3 | 0.06 | 1.18 | 1.12 | 66 | 0.22 | 0.1 | 0.2 | 8.02 | 25.8 | 0.25 |
| S_DS1C2 | Mar | 22 | 2.6 | 0.09 | 1.77 | 1.68 | 74 | 0.29 | 0.03 | 0.2 | 7.96 | 24.4 | 0.25 |
| S_DS1C3 | Apr | 20 | 3.4 | 0.01 | 1.69 | 1.68 | 72 | 0.37 | 0.03 | 0.1 | 8.38 | 21.2 | 0.3 |
| S_DS1C4 | May | 20 | 3 | 0.07 | 1.74 | 1.67 | 85 | 0.52 | 0.03 | 0.35 | 8.04 | 16.6 | 0.33 |
| S_DS1C5 | Jun | 19 | 3.5 | 0.11 | 1.79 | 1.68 | 78 | 0.45 | 0.03 | 0.18 | 8.38 | 15.2 | 0.18 |
| S_DS1C6 | Jul | 12 |  | 0.64 | 1.97 | 1.33 |  | 0.48 | 0.03 | 0.83 | 8.01 | 13.2 | 0.14 |
| S_DS1C7 | Aug | 17 |  | 0.41 | 2.01 | 1.6 |  | 0.48 | 0.03 | 0.36 | 8.16 | 12.3 | 0.24 |
| S_DS1C8 | Sept | 17 |  | 0.2 | 2.07 | 1.87 |  | 0.5 | 0.03 | 0.38 | 7.93 | 14.7 | 0.27 |
| S_DS2A1 | Feb | 19 | 3.8 | 0.09 | 1.15 | 1.06 | 70 | 0.17 | 0.16 | 0.18 | 8.06 | 25.3 | 0.25 |
| S_DS2A2 | Mar | 24 | 2.7 | 0.05 | 1.24 | 1.19 | 73 | 0.24 | 0.03 | 0.16 | 7.96 | 23.5 | 0.25 |
| S_DS2A3 | Apr | 20 | 3.4 | 0.1 | 1.5 | 1.4 | 71 | 0.31 | 0.04 | 0.28 | 8.02 | 22.2 | 0.28 |
| S_DS2A4 | May | 21 | 3.4 | 0.09 | 1.45 | 1.36 | 86 | 0.43 | 0.07 | 0.2 | 8.05 | 17.1 | 0.39 |
| S_DS2A5 | Jun | 20 | 2.3 | 0.08 | 1.29 | 1.21 | 83 | 0.45 | 0.03 | 0.18 | 8.18 | 18 | 0.33 |
| S_DS2A6 | Jul | 11 |  | 0.88 | 1.89 | 1.01 |  | 0.45 | 0.03 | 0.62 | 8.14 | 13.2 | 0.17 |
| S_DS2A7 | Aug | 17 |  | 0.07 | 1.87 | 1.8 |  | 0.43 | 0.03 | 0.27 | 8.2 | 11.8 | 0.22 |
| S_DS2A8 | Sept | 17 |  | 0.14 | 1.85 | 1.71 |  | 0.5 | 0.03 | 0.33 | 7.95 | 15.1 | 0.28 |
| S_DS2B1 | Feb | 19 | 3.2 | 0.09 | 0.91 | 0.82 | 71 | 0.16 | 0.18 | 0.28 | 8.11 | 25 | 0.26 |
| S_DS2B2 | Mar | 22 | 2.7 | 0.03 | 1.41 | 1.38 | 73 | 0.27 | 0.03 | 0.16 | 7.99 | 24.1 | 0.27 |
| S_DS2B3 | Apr | 20 | 3.4 | 0.08 | 1.5 | 1.42 | 70 | 0.32 | 0.03 | 0.16 | 8.22 | 22.2 | 0.29 |
| S_DS2B4 | May | 20 | 3.3 | 0.05 | 1.47 | 1.42 | 85 | 0.44 | 0.05 | 0.11 | 8.04 | 16.4 |  |
| S_DS2B5 | Jun | 20 | 2.3 | 0.06 | 1.67 | 1.61 | 84 | 0.42 | 0.03 | 0.2 | 8.27 | 16.7 | 0.3 |
| S_DS2B6 | Jul | 12 |  | 0.5 | 1.92 | 1.42 |  | 0.47 | 0.03 | 0.26 | 8.18 | 13.1 | 0.25 |
| S_DS2B7 | Aug | 17 |  | 0.08 | 1.98 | 1.9 |  | 0.45 | 0.03 | 0.27 | 8.24 | 11.7 | 0.23 |
| S_DS2B8 | Sept | 17 |  | 0.14 | 1.79 | 1.65 |  | 0.5 | 0.03 | 0.42 | 7.95 | 14.8 | 0.21 |
| S_DS2C1 | Feb | 18 | 3.3 | 0.03 | 0.93 | 0.9 | 66 | 0.11 | 0.16 | 0.23 | 7.92 | 24.4 | 0.28 |
| S_DS2C2 | Mar | 22 | 2.7 | 0.05 | 1.53 | 1.48 | 74 | 0.25 | 0.03 | 0.22 | 8.01 | 24.6 | 0.29 |
| S_DS2C3 | Apr | 20 | 3.2 | 0.06 | 1.49 | 1.43 | 69 | 0.32 | 0.03 | 0.1 | 7.83 | 22 | 0.31 |
| S_DS2C4 | May | 21 | 3.2 | 0.04 | 1.64 | 1.6 | 81 | 0.5 | 0.03 | 0.35 | 8.37 | 16.3 | 0.36 |
| S_DS2C5 | Jun | 19 | 2.5 | 0.04 | 1.53 | 1.49 | 77 | 0.43 | 0.03 | 0.18 | 8.36 | 16.4 | 0.39 |
| S_DS2C6 | Jul | 12 |  | 0.37 | 1.99 | 1.62 |  | 0.45 | 0.03 | 0.21 | 8.28 | 13.2 | 0.26 |
| S_DS2C7 | Aug | 17 |  | 0.42 | 2.05 | 1.63 |  | 0.52 | 0.03 | 0.27 | 8.23 | 11.8 | 0.22 |
| S_DS2C8 | Sept | 17 |  | 0.1 | 1.92 | 1.82 |  | 0.52 | 0.03 | 0.25 | 7.95 | 15.1 | 0.23 |

**Table S3:** The number of samples collected and processed

|  | **Number of samples collected** | | **Number of samples successfully sequenced** | |
| --- | --- | --- | --- | --- |
| **Sample** | **System R** | **System S** | **System R** | **System S** |
| **SW** | 8 | 8 | 7 | 6 |
| **FI** | 8 | 8 | 8 | 8 |
| **FB** | 8 | 8 | 8 | 8 |
| **FE** | 8 | 8 | 8 | 8 |
| **CHL** | 16 | 16 | 12 | 9 |
| **CHM** | 8 | 16 | 5 | 11 |
| **DS1** | 16 | 24 | 11 | 23 |
| **DS2** | 24 | 24 | 19 | 21 |

**Table S4**: Mean relative abundance of dominant bacterial phyla (> 0.1%) across all samples within the DWTPs and corresponding DWDS lines of both systems. Values depicted as *Mean* ± *SD* for each location within each system

| **System R** | | | | | | | | | |
| --- | --- | --- | --- | --- | --- | --- | --- | --- | --- |
| **Phyla** | **R_SW** | **R_FI** | **R_FB** | **F_FE** | **R_CHLA** | **R_CHLB** | **R_CHM** | **R_DS1** | **R_DS2** |
| *Proteobacteria* | 0.42 ± 0.07 | 0.23 ± 0.02 | 0.35 ± 0.1 | 0.3 ± 0.15 | 0.11 ± 0.09 | 0.16 ± 0.11 | 0.49 ± 0.28 | 0.56 ± 0.26 | 0.69 ± 0.18 |
| *Alphaproteobacteria* | 0.05 ± 0.02 | 0.05 ± 0.02 | 0.12 ± 0.07 | 0.12 ± 0.15 | 0.05 ± 0.03 | 0.05 ± 0.05 | 0.16 ± 0.15 | 0.13 ± 0.09 | 0.3 ± 0.18 |
| *Gammaproteobacteria*  *Betaproteobacteriales* | 0.22 ± 0.04 | 0.12 ± 0.03 | 0.11 ± 0.05 | 0.12 ± 0.05 | 0.04 ± 0.07 | 0.07 ± 0.06 | 0.07 ± 0.04 | 0.22 ± 0.22 | 0.25 ± 0.16 |
| *Gammaproteobacteria*  Other | 0.13 ± 0.08 | 0.05 ± 0.02 | 0.12 ± 0.06 | 0.05 ± 0.04 | 0.03 ± 0.02 | 0.04 ± 0.05 | 0.25 ± 0.29 | 0.21 ± 0.32 | 0.13 ± 0.15 |
| *Deltaproteobacteria* | 0.01 ± 0 | 0 ± 0 | 0 ± 0 | 0.01 ± 0.01 | 0 ± 0 | 0 ± 0 | 0.01 ± 0.02 | 0 ± 0 | 0.01 ± 0.01 |
| *Epsilonproteobacteria* | 0 ± 0 | 0 ± 0 | 0 ± 0 | 0 ± 0 | 0 ± 0 | 0 ± 0 | 0 ± 0 | 0 ± 0 | 0 ± 0 |
| Unclassified *Proteobacteria* | 0 ± 0 | 0 ± 0 | 0 ± 0 | 0 ± 0 | 0 ± 0 | 0 ± 0 | 0 ± 0.01 | 0 ± 0 | 0 ± 0 |
| *Actinobacteria* | 0.21 ± 0.07 | 0.31 ± 0.06 | 0.2 ± 0.07 | 0.28 ± 0.1 | 0.06 ± 0.13 | 0.09 ± 0.1 | 0.03 ± 0.02 | 0.04 ± 0.05 | 0.02 ± 0.04 |
| *Bacteroidetes* | 0.13 ± 0.09 | 0.17 ± 0.05 | 0.2 ± 0.03 | 0.14 ± 0.08 | 0.03 ± 0.06 | 0.12 ± 0.19 | 0.02 ± 0.03 | 0.04 ± 0.09 | 0.03 ± 0.03 |
| *Planctomycetes* | 0.06 ± 0.07 | 0.02 ± 0.02 | 0.03 ± 0.01 | 0.03 ± 0.03 | 0.18 ± 0.18 | 0.03 ± 0.06 | 0.25 ± 0.24 | 0.17 ± 0.15 | 0.14 ± 0.1 |
| *Acidobacteria* | 0.03 ± 0.02 | 0.12 ± 0.03 | 0.06 ± 0.03 | 0.13 ± 0.05 | 0.02 ± 0.04 | 0.01 ± 0.03 | 0.01 ± 0.02 | 0.02 ± 0.02 | 0.02 ± 0.02 |
| *Cyanobacteria* | 0.05 ± 0.03 | 0.02 ± 0.01 | 0.05 ± 0.03 | 0.01 ± 0 | 0.02 ± 0.02 | 0.1 ± 0.11 | 0.03 ± 0.02 | 0.02 ± 0.02 | 0.01 ± 0.01 |
| *Verrucomicrobia* | 0.03 ± 0.02 | 0.03 ± 0.01 | 0.03 ± 0.02 | 0.04 ± 0.02 | 0 ± 0.01 | 0.01 ± 0.02 | 0.02 ± 0.04 | 0.02 ± 0.02 | 0.01 ± 0.01 |
| *Firmicutes* | 0 ± 0 | 0 ± 0.01 | 0 ± 0 | 0 ± 0 | 0.01 ± 0.01 | 0.01 ± 0.02 | 0.01 ± 0.01 | 0.01 ± 0.02 | 0.01 ± 0.02 |
| *Nitrospirae* | 0 ± 0 | 0.01 ± 0 | 0 ± 0 | 0.01 ± 0.01 | 0 ± 0 | 0 ± 0 | 0 ± 0.01 | 0.01 ± 0.01 | 0.01 ± 0.01 |
| *Chlorobi* | 0.01 ± 0.01 | 0.01 ± 0.01 | 0.01 ± 0.01 | 0.01 ± 0 | 0 ± 0 | 0 ± 0.01 | 0 ± 0 | 0 ± 0 | 0 ± 0 |
| *Chlamydiae* | 0 ± 0 | 0.01 ± 0.01 | 0 ± 0 | 0.01 ± 0.01 | 0 ± 0 | 0 ± 0 | 0 ± 0 | 0 ± 0.01 | 0 ± 0 |
| *Gemmatimonadetes* | 0 ± 0 | 0.01 ± 0 | 0 ± 0 | 0.01 ± 0 | 0 ± 0 | 0 ± 0 | 0 ± 0 | 0.01 ± 0.01 | 0 ± 0 |
| *Chloroflexi* | 0 ± 0 | 0.01 ± 0.03 | 0 ± 0 | 0 ± 0 | 0 ± 0 | 0 ± 0.01 | 0 ± 0 | 0 ± 0 | 0 ± 0 |
| *Elusimicrobia* | 0 ± 0 | 0 ± 0 | 0 ± 0 | 0 ± 0 | 0 ± 0 | 0 ± 0 | 0 ± 0 | 0.01 ± 0.02 | 0.01 ± 0.01 |
| *Ignavibacteriae* | 0 ± 0 | 0 ± 0 | 0 ± 0 | 0 ± 0 | 0 ± 0 | 0 ± 0 | 0 ± 0 | 0.01 ± 0.01 | 0.01 ± 0.01 |
| *Parcubacteria* | 0 ± 0 | 0 ± 0 | 0 ± 0 | 0 ± 0 | 0 ± 0 | 0 ± 0 | 0 ± 0 | 0 ± 0 | 0 ± 0 |
| *Armatimonadetes* | 0 ± 0 | 0 ± 0 | 0 ± 0 | 0 ± 0 | 0 ± 0 | 0 ± 0 | 0 ± 0 | 0 ± 0 | 0 ± 0 |
| Unclassified | 0 ± 0 | 0 ± 0 | 0.01 ± 0 | 0 ± 0 | 0.15 ± 0.07 | 0.07 ± 0.05 | 0.02 ± 0.02 | 0.01 ± 0.02 | 0.01 ± 0.01 |
| Remaining 33 Phyla (<0.1%) | 0.01 ± 0.01 | 0 ± 0 | 0 ± 0 | 0 ± 0 | 0 ± 0 | 0.01 ± 0.02 | 0 ± 0 | 0 ± 0 | 0 ± 0.01 |

| **System S** | | | | | | | | | |
| --- | --- | --- | --- | --- | --- | --- | --- | --- | --- |
| **Phyla** | **S_SW** | **S_FI** | **S_FB** | **S_FE** | **S_CHLA** | **S_CHLB** | **S_CHM** | **S_DS1** | **S_DS2** |
| *Proteobacteria* | 0.24 ± 0.09 | 0.27 ± 0.04 | 0.42 ± 0.12 | 0.26 ± 0.1 | 0.3 ± 0.19 | 0.26 ± 0.2 | 0.35 ± 0.24 | 0.76 ± 0.17 | 0.72 ± 0.18 |
| *Alphaproteobacteria* | 0.05 ± 0.01 | 0.04 ± 0.01 | 0.22 ± 0.18 | 0.05 ± 0.02 | 0.21 ± 0.14 | 0.17 ± 0.21 | 0.2 ± 0.18 | 0.49 ± 0.21 | 0.43 ± 0.19 |
| *Gammaproteobacteria*  *Betaproteobacteriales* | 0.12 ± 0.04 | 0.18 ± 0.04 | 0.14 ± 0.07 | 0.18 ± 0.09 | 0.05 ± 0.06 | 0.02 ± 0.01 | 0.04 ± 0.03 | 0.18 ± 0.12 | 0.24 ± 0.15 |
| *Gammaproteobacteria*  Other | 0.07 ± 0.05 | 0.04 ± 0.02 | 0.05 ± 0.03 | 0.03 ± 0.01 | 0.03 ± 0.03 | 0.06 ± 0.07 | 0.09 ± 0.15 | 0.06 ± 0.13 | 0.04 ± 0.1 |
| *Deltaproteobacteria* | 0.01 ± 0 | 0 ± 0 | 0.01 ± 0.01 | 0 ± 0 | 0.01 ± 0.01 | 0.01 ± 0.01 | 0.02 ± 0.02 | 0.02 ± 0.04 | 0.01 ± 0.01 |
| *Epsilonproteobacteria* | 0 ± 0 | 0 ± 0 | 0 ± 0 | 0 ± 0 | 0 ± 0 | 0 ± 0 | 0 ± 0 | 0 ± 0 | 0 ± 0 |
| Unclassified *Proteobacteria* | 0 ± 0 | 0 ± 0 | 0 ± 0 | 0 ± 0 | 0 ± 0 | 0 ± 0 | 0 ± 0 | 0 ± 0 | 0 ± 0 |
| *Actinobacteria* | 0.31 ± 0.02 | 0.31 ± 0.08 | 0.18 ± 0.05 | 0.31 ± 0.07 | 0.06 ± 0.06 | 0.01 ± 0.01 | 0.03 ± 0.03 | 0.02 ± 0.01 | 0.03 ± 0.05 |
| *Bacteroidetes* | 0.09 ± 0.03 | 0.18 ± 0.09 | 0.18 ± 0.09 | 0.14 ± 0.04 | 0.01 ± 0.01 | 0.01 ± 0.01 | 0.06 ± 0.09 | 0.03 ± 0.03 | 0.03 ± 0.03 |
| *Planctomycetes* | 0.04 ± 0.02 | 0.01 ± 0 | 0.02 ± 0.01 | 0.01 ± 0.01 | 0.08 ± 0.08 | 0.04 ± 0.07 | 0.25 ± 0.16 | 0.08 ± 0.07 | 0.08 ± 0.08 |
| *Acidobacteria* | 0.06 ± 0.01 | 0.12 ± 0.03 | 0.05 ± 0.02 | 0.15 ± 0.03 | 0.01 ± 0.01 | 0 ± 0 | 0.01 ± 0.01 | 0.01 ± 0.01 | 0.02 ± 0.02 |
| *Cyanobacteria* | 0.04 ± 0.02 | 0.02 ± 0.01 | 0.04 ± 0.02 | 0.01 ± 0.01 | 0.08 ± 0.04 | 0.11 ± 0.1 | 0.08 ± 0.05 | 0.03 ± 0.03 | 0.03 ± 0.03 |
| *Verrucomicrobia* | 0.03 ± 0.01 | 0.02 ± 0 | 0.01 ± 0.01 | 0.02 ± 0.01 | 0 ± 0 | 0 ± 0 | 0 ± 0 | 0 ± 0 | 0 ± 0 |
| *Firmicutes* | 0 ± 0 | 0 ± 0 | 0 ± 0 | 0 ± 0 | 0.02 ± 0.02 | 0.03 ± 0.05 | 0.03 ± 0.03 | 0.01 ± 0.01 | 0.01 ± 0.01 |
| *Nitrospirae* | 0.01 ± 0 | 0.01 ± 0.01 | 0.01 ± 0.01 | 0.02 ± 0.01 | 0 ± 0.01 | 0 ± 0 | 0 ± 0 | 0.01 ± 0 | 0.01 ± 0.01 |
| *Chlorobi* | 0.02 ± 0.01 | 0.01 ± 0.01 | 0.01 ± 0.01 | 0.01 ± 0.01 | 0 ± 0 | 0 ± 0 | 0 ± 0 | 0 ± 0 | 0 ± 0 |
| *Chlamydiae* | 0.01 ± 0 | 0.01 ± 0.01 | 0.01 ± 0.01 | 0.01 ± 0.01 | 0 ± 0 | 0 ± 0 | 0 ± 0 | 0 ± 0 | 0 ± 0 |
| *Gemmatimonadetes* | 0 ± 0 | 0 ± 0 | 0 ± 0 | 0 ± 0 | 0 ± 0 | 0 ± 0 | 0 ± 0 | 0 ± 0 | 0 ± 0 |
| *Chloroflexi* | 0.01 ± 0 | 0 ± 0 | 0 ± 0 | 0 ± 0 | 0 ± 0 | 0 ± 0 | 0 ± 0 | 0 ± 0.01 | 0 ± 0 |
| *Elusimicrobia* | 0 ± 0 | 0 ± 0 | 0 ± 0 | 0 ± 0 | 0 ± 0 | 0 ± 0 | 0 ± 0 | 0 ± 0 | 0.02 ± 0.03 |
| *Ignavibacteriae* | 0.01 ± 0.01 | 0 ± 0 | 0 ± 0 | 0 ± 0 | 0 ± 0 | 0 ± 0 | 0 ± 0 | 0.01 ± 0.04 | 0 ± 0 |
| *Parcubacteria* | 0.02 ± 0.02 | 0 ± 0 | 0 ± 0 | 0 ± 0 | 0 ± 0 | 0 ± 0 | 0 ± 0 | 0 ± 0 | 0 ± 0 |
| *Armatimonadetes* | 0 ± 0 | 0 ± 0 | 0 ± 0 | 0 ± 0 | 0 ± 0 | 0 ± 0 | 0 ± 0 | 0 ± 0 | 0 ± 0 |
| Unclassified | 0.02 ± 0.02 | 0 ± 0 | 0 ± 0 | 0 ± 0 | 0.11 ± 0.09 | 0.16 ± 0.05 | 0.02 ± 0.02 | 0.01 ± 0.01 | 0.01 ± 0.01 |
| Remaining 33 Phyla (<0.1%) | 0.02 ± 0.02 | 0 ± 0 | 0.01 ± 0.01 | 0 ± 0 | 0.01 ± 0.01 | 0 ± 0.01 | 0 ± 0 | 0 ± 0 | 0 ± 0.01 |

*Mean ± Standard deviation

**Table S5**: Mean and standard deviations of the number of sequences and alpha diversity indexes averaged over the duration of the study for each individual study site

|  | **Number of sequences** | | **Number of observed taxa (Sobs)** | | **Inverse Simpson Diversity Index** | | **Shannon Diversity Index** | | **Pielou's evenness** | | **Good's coverage** | |
| --- | --- | --- | --- | --- | --- | --- | --- | --- | --- | --- | --- | --- |
|  | ***Mean*** | ***SD*** | ***Mean*** | ***SD*** | ***Mean*** | ***SD*** | ***Mean*** | ***SD*** | ***Mean*** | ***SD*** | ***Mean*** | ***SD*** |
| R_SW | 34465.71 | 19422.11 | 208.52 | 70.03 | 39.58 | 16.05 | 4.30 | 0.55 | 0.82 | 0.03 | 0.94 | 0.03 |
| S_SW | 60236.00 | 17308.25 | 304.32 | 17.09 | 40.69 | 5.40 | 4.66 | 0.10 | 0.82 | 0.01 | 0.88 | 0.01 |
| R_FI | 35137.71 | 15165.55 | 196.28 | 52.47 | 36.04 | 11.11 | 4.27 | 0.30 | 0.81 | 0.02 | 0.94 | 0.02 |
| S_FI | 49732.13 | 13430.06 | 185.92 | 28.20 | 27.60 | 8.75 | 4.04 | 0.31 | 0.77 | 0.04 | 0.94 | 0.01 |
| R_FB | 39244.88 | 21034.37 | 229.43 | 70.23 | 43.79 | 19.38 | 4.39 | 0.62 | 0.81 | 0.06 | 0.92 | 0.03 |
| S_FB | 46060.38 | 28383.14 | 232.39 | 56.01 | 42.36 | 13.16 | 4.41 | 0.33 | 0.81 | 0.03 | 0.92 | 0.03 |
| R_FE | 35808.13 | 13983.04 | 159.10 | 32.14 | 28.98 | 9.96 | 4.01 | 0.41 | 0.79 | 0.05 | 0.96 | 0.01 |
| S_FE | 51383.50 | 24773.13 | 160.79 | 20.54 | 21.84 | 6.60 | 3.86 | 0.20 | 0.76 | 0.04 | 0.95 | 0.01 |
| R_CHLA | 18781.29 | 16926.91 | 58.35 | 42.94 | 9.94 | 9.42 | 2.64 | 0.67 | 0.68 | 0.07 | 0.99 | 0.01 |
| S_CHLA | 15748.00 | 11454.43 | 82.79 | 36.36 | 13.08 | 7.56 | 3.14 | 0.55 | 0.72 | 0.06 | 0.99 | 0.01 |
| R_CHLB | 18146.60 | 28495.57 | 80.63 | 76.63 | 15.65 | 10.93 | 3.02 | 0.97 | 0.75 | 0.09 | 0.98 | 0.03 |
| S_CHLB | 15234.00 | 25980.55 | 48.23 | 25.33 | 8.86 | 2.17 | 2.70 | 0.17 | 0.72 | 0.07 | 0.99 | 0.01 |
| R_CHM | 10565.40 | 6432.02 | 78.96 | 31.22 | 12.81 | 9.36 | 2.98 | 0.83 | 0.69 | 0.15 | 0.99 | 0.01 |
| S_CHM | 18030.45 | 14458.32 | 86.94 | 33.26 | 12.77 | 6.01 | 3.14 | 0.44 | 0.71 | 0.07 | 0.98 | 0.01 |
| R_DS1 | 29708.09 | 25598.37 | 106.48 | 41.71 | 18.13 | 10.94 | 3.26 | 1.04 | 0.70 | 0.19 | 0.98 | 0.01 |
| S_DS1 | 23499.52 | 14113.34 | 99.37 | 47.07 | 10.37 | 7.19 | 2.87 | 0.75 | 0.63 | 0.12 | 0.97 | 0.02 |
| R_DS2 | 25236.84 | 17138.83 | 129.81 | 37.40 | 16.04 | 7.88 | 3.50 | 0.52 | 0.72 | 0.08 | 0.97 | 0.02 |
| S_DS2 | 26578.38 | 16234.82 | 115.24 | 45.57 | 12.99 | 8.37 | 3.14 | 0.80 | 0.66 | 0.12 | 0.97 | 0.02 |

**Table S6**: Mean percentage relative abundance of the top 33 abundant sequence variants (MRA > 0.5%) across all samples from both systems

| **System R** | | | | | | | | | | |
| --- | --- | --- | --- | --- | --- | --- | --- | --- | --- | --- |
| **ASV** | **ASV Taxonomy** | **R_RW** | **R_FI** | **R_FB** | **R_FE** | **R_CHLA** | **R_CHLB** | **R_CHM** | **R_DS1** | **R_DS2** |
| ASV_1 | *Proteobacteria_Alphaproteobacteria_Rhizobiales_Beijerinckiaceae_Methylobacterium* | 0.00 | 0.00 | 0.00 | 0.00 | 0.00 | 0.63 | 2.36 | 2.82 | 9.14 |
| ASV_2 | *Actinobacteria_Actinobacteria_Frankiales_Sporichthyaceae* | 4.82 | 9.48 | 6.11 | 8.43 | 2.29 | 2.82 | 0.38 | 0.65 | 0.33 |
| ASV_3 | *Planctomycetes_Planctomycetacia_Gemmatales_Gemmataceae* | 0.45 | 0.38 | 0.22 | 0.94 | 14.20 | 1.17 | 10.63 | 8.64 | 6.72 |
| ASV_4 | *Acidobacteria_Holophagae_Holophagales_Holophagaceae* | 2.01 | 5.62 | 3.30 | 7.14 | 0.89 | 0.64 | 0.79 | 0.62 | 0.56 |
| ASV_5 | *Actinobacteria_Actinobacteria_Frankiales_Sporichthyaceae* | 5.37 | 5.15 | 4.38 | 4.67 | 1.00 | 0.90 | 0.42 | 0.43 | 0.14 |
| ASV_6 | *Proteobacteria_Gammaproteobacteria_Betaproteobacteriales_Nitrosomonadaceae_Nitrosomonas* | 0.01 | 0.00 | 0.01 | 0.00 | 0.00 | 0.00 | 0.49 | 1.45 | 10.92 |
| ASV_7 | *Actinobacteria_Actinobacteria_Frankiales_Sporichthyaceae* | 2.95 | 5.22 | 2.23 | 4.75 | 0.93 | 1.77 | 0.24 | 0.60 | 0.23 |
| ASV_8 | *Proteobacteria_Gammaproteobacteria_Betaproteobacteriales_Burkholderiaceae_Limnohabitans* | 2.18 | 1.76 | 1.02 | 2.74 | 0.45 | 1.45 | 0.77 | 0.07 | 0.00 |
| ASV_10 | *Proteobacteria_Gammaproteobacteria_Pseudomonadales_Pseudomonadaceae_Pseudomonas* | 0.52 | 2.69 | 4.45 | 2.69 | 0.48 | 0.87 | 3.61 | 0.94 | 1.69 |
| ASV_11 | *Proteobacteria_Alphaproteobacteria_Sphingomonadales_Sphingomonadaceae_Sphingomonas* | 0.00 | 0.06 | 3.91 | 3.89 | 0.48 | 0.11 | 2.50 | 1.59 | 4.92 |
| ASV_12 | *Acidobacteria* | 0.62 | 3.25 | 1.50 | 3.79 | 0.94 | 0.59 | 0.02 | 0.18 | 0.08 |
| ASV_13 | *Proteobacteria_Alphaproteobacteria_Rhizobiales_Phreatobacter* | 0.00 | 0.00 | 0.00 | 0.00 | 0.00 | 0.00 | 0.10 | 0.60 | 1.68 |
| ASV_15 | *Thaumarchaeota_Nitrososphaeria_ Nitrosopumilales_Nitrosopumilaceae_Candidatus Nitrosoarchaeum* | 2.76 | 0.98 | 1.47 | 0.99 | 0.07 | 0.00 | 0.18 | 0.09 | 0.02 |
| ASV_16 | *Proteobacteria_Gammaproteobacteria_Pseudomonadales_Pseudomonadaceae_Pseudomonas* | 0.68 | 0.07 | 0.10 | 0.00 | 0.00 | 0.00 | 0.00 | 6.67 | 0.00 |
| ASV_17 | *Actinobacteria_Acidimicrobiia_Acidimicrobiales_Acidimicrobiaceae* | 0.65 | 2.65 | 0.75 | 2.77 | 0.43 | 1.17 | 0.08 | 0.30 | 0.15 |
| ASV_18 | *Bacteroidetes_Flavobacteriia_Flavobacteriales_Flavobacteriaceae* | 0.00 | 0.35 | 0.92 | 0.15 | 0.02 | 0.54 | 0.00 | 0.02 | 0.06 |
| ASV_19 | *Proteobacteria_Gammaproteobacteria_Pseudomonadales_Moraxellaceae_Acinetobacter* | 0.00 | 0.00 | 0.01 | 0.00 | 0.56 | 0.14 | 6.66 | 5.00 | 0.16 |
| ASV_22 | *Proteobacteria_Gammaproteobacteria_Betaproteobacteriales_Burkholderiaceae* | 1.87 | 1.00 | 1.32 | 0.81 | 0.25 | 0.00 | 0.00 | 0.04 | 0.66 |
| ASV_23 | *Proteobacteria_Gammaproteobacteria_Beggiatoales_Beggiatoaceae* | 0.00 | 0.00 | 0.00 | 0.00 | 0.00 | 0.00 | 0.00 | 0.40 | 4.94 |
| ASV_25 | *Proteobacteria_Alphaproteobacteria_Rhodobacterales_Rhodobacteraceae_Cereibacter* | 0.00 | 0.01 | 0.29 | 0.08 | 0.00 | 0.00 | 0.39 | 0.72 | 0.30 |
| ASV_26 | *Cyanobacteria_Melainabacteria_Obscuribacterales* | 0.00 | 0.00 | 0.00 | 0.01 | 0.15 | 0.26 | 0.82 | 0.66 | 0.30 |
| ASV_29 | *Proteobacteria_Alphaproteobacteria_Caulobacterales_Hyphomonadaceae* | 0.00 | 0.00 | 0.00 | 0.00 | 0.00 | 0.00 | 2.94 | 0.41 | 0.19 |
| ASV_30 | *Planctomycetes_Planctomycetacia_Planctomycetales* | 0.00 | 0.00 | 0.11 | 0.10 | 0.63 | 0.00 | 4.59 | 3.18 | 2.61 |
| ASV_31 | *Proteobacteria_Gammaproteobacteria_Pseudomonadales_Moraxellaceae_Acinetobacter* | 0.00 | 0.00 | 0.00 | 0.00 | 0.00 | 0.00 | 5.15 | 3.86 | 0.06 |
| ASV_32 | *Proteobacteria_Alphaproteobacteria_Rhizobiales_Hyphomicrobiaceae_Hyphomicrobium* | 0.00 | 0.00 | 0.00 | 0.00 | 0.00 | 0.00 | 0.05 | 0.68 | 1.07 |
| ASV_33 | *Proteobacteria_Alphaproteobacteria_Sphingomonadales_Sphingomonadaceae_Sphingomonas* | 0.00 | 0.00 | 0.00 | 0.00 | 0.00 | 0.00 | 0.00 | 0.37 | 1.45 |
| ASV_34 | *Proteobacteria_Gammaproteobacteria_Betaproteobacteriales_Gallionellaceae* | 0.00 | 0.00 | 0.00 | 0.00 | 0.00 | 0.00 | 0.00 | 1.78 | 2.56 |
| ASV_36 | *Planctomycetes_Phycisphaerae_Phycisphaerales_Phycisphaeraceae* | 0.00 | 0.00 | 0.00 | 0.01 | 0.65 | 0.08 | 2.85 | 0.56 | 0.58 |
| ASV_37 | *Proteobacteria_Gammaproteobacteria_Betaproteobacteriales_Sulfuricellaceae_Sulfuricella* | 0.00 | 0.00 | 0.00 | 0.00 | 0.00 | 0.00 | 0.01 | 0.99 | 1.30 |
| ASV_40 | *Proteobacteria_Alphaproteobacteria_Rickettsiales* | 0.00 | 0.03 | 0.05 | 0.01 | 5.20 | 2.27 | 0.57 | 0.19 | 0.10 |
| ASV_41 | *Proteobacteria_Alphaproteobacteria_Rickettsiales* | 0.01 | 0.00 | 0.00 | 0.01 | 6.71 | 3.36 | 0.81 | 0.80 | 0.26 |
| ASV_44 | *Bacteroidetes_Flavobacteriia_Flavobacteriales_Flavobacteriaceae_Flavobacterium* | 4.13 | 1.64 | 3.64 | 0.35 | 0.07 | 0.39 | 0.01 | 0.00 | 0.02 |
| ASV_80 | *Proteobacteria_Alphaproteobacteria_Rhizobiales_Beijerinckiaceae_Methylobacterium* | 0.00 | 0.00 | 0.00 | 0.00 | 0.46 | 0.05 | 0.00 | 0.00 | 0.01 |
| **System S** | | | | | | | | | | |
| **ASV** | **ASV Taxonomy** | **S_SW** | **S_FI** | **S_FB** | **S_FE** | **S_CHLA** | **S_CHLB** | **S_CHM** | **S_DS1** | **S_DS2** |
| ASV_1 | *Proteobacteria_Alphaproteobacteria_Rhizobiales_Beijerinckiaceae_Methylobacterium* | 0.01 | 0.01 | 0.00 | 0.00 | 0.91 | 0.06 | 0.74 | 19.31 | 15.19 |
| ASV_2 | *Actinobacteria_Actinobacteria_Frankiales_Sporichthyaceae* | 8.42 | 9.75 | 5.74 | 9.55 | 1.47 | 0.56 | 0.48 | 0.36 | 0.86 |
| ASV_3 | *Planctomycetes_Planctomycetacia_Gemmatales_Gemmataceae* | 0.41 | 0.19 | 0.11 | 0.25 | 4.20 | 1.24 | 14.92 | 4.68 | 4.83 |
| ASV_4 | *Acidobacteria_Holophagae_Holophagales_Holophagaceae* | 1.69 | 7.14 | 2.32 | 9.44 | 0.87 | 0.01 | 0.39 | 0.55 | 0.94 |
| ASV_5 | *Actinobacteria_Actinobacteria_Frankiales_Sporichthyaceae* | 7.63 | 6.34 | 4.45 | 5.91 | 1.43 | 0.05 | 0.27 | 0.16 | 0.45 |
| ASV_6 | *Proteobacteria_Gammaproteobacteria_Betaproteobacteriales_Nitrosomonadaceae_Nitrosomonas* | 0.01 | 0.04 | 0.02 | 0.03 | 1.10 | 0.41 | 0.36 | 10.32 | 12.96 |
| ASV_7 | *Actinobacteria_Actinobacteria_Frankiales_Sporichthyaceae* | 4.28 | 6.25 | 2.25 | 5.72 | 1.20 | 0.04 | 0.22 | 0.23 | 0.40 |
| ASV_8 | *Proteobacteria_Gammaproteobacteria_Betaproteobacteriales_Burkholderiaceae_Limnohabitans* | 0.41 | 7.26 | 2.49 | 10.47 | 0.64 | 0.03 | 0.12 | 0.03 | 0.30 |
| ASV_10 | *Proteobacteria_Gammaproteobacteria_Pseudomonadales_Pseudomonadaceae_Pseudomonas* | 0.32 | 0.85 | 1.04 | 0.46 | 0.09 | 0.00 | 0.14 | 1.36 | 1.63 |
| ASV_11 | *Proteobacteria_Alphaproteobacteria_Sphingomonadales_Sphingomonadaceae_Sphingomonas* | 0.05 | 0.03 | 2.32 | 0.28 | 0.92 | 0.06 | 1.43 | 3.61 | 4.70 |
| ASV_12 | *Acidobacteria* | 1.60 | 3.40 | 1.29 | 4.13 | 0.13 | 0.02 | 0.05 | 0.22 | 0.30 |
| ASV_13 | *Proteobacteria_Alphaproteobacteria_Rhizobiales_Phreatobacter* | 0.00 | 0.00 | 0.00 | 0.00 | 2.48 | 0.19 | 0.14 | 7.05 | 7.92 |
| ASV_15 | *Thaumarchaeota_Nitrososphaeria_ Nitrosopumilales_Nitrosopumilaceae_Candidatus Nitrosoarchaeum* | 6.40 | 1.00 | 2.08 | 0.79 | 0.11 | 0.01 | 0.02 | 0.04 | 0.07 |
| ASV_16 | *Proteobacteria_Gammaproteobacteria_Pseudomonadales_Pseudomonadaceae_Pseudomonas* | 0.01 | 0.00 | 0.00 | 0.00 | 0.00 | 3.18 | 0.00 | 0.00 | 0.00 |
| ASV_17 | *Actinobacteria_Acidimicrobiia_Acidimicrobiales_Acidimicrobiaceae* | 0.93 | 1.56 | 0.70 | 2.86 | 0.14 | 0.00 | 0.03 | 0.09 | 0.19 |
| ASV_18 | *Bacteroidetes_Flavobacteriia_Flavobacteriales_Flavobacteriaceae* | 0.13 | 2.23 | 5.54 | 1.25 | 0.09 | 0.00 | 0.03 | 0.06 | 0.06 |
| ASV_19 | *Proteobacteria_Gammaproteobacteria_Pseudomonadales_Moraxellaceae_Acinetobacter* | 0.02 | 0.00 | 0.00 | 0.00 | 0.03 | 0.40 | 0.31 | 0.24 | 0.12 |
| ASV_22 | *Proteobacteria_Gammaproteobacteria_Betaproteobacteriales_Burkholderiaceae* | 0.06 | 1.91 | 1.17 | 0.63 | 0.00 | 0.00 | 0.10 | 0.05 | 0.45 |
| ASV_23 | *Proteobacteria_Gammaproteobacteria_Beggiatoales_Beggiatoaceae* | 0.00 | 0.00 | 0.00 | 0.00 | 0.00 | 0.00 | 0.00 | 0.19 | 0.01 |
| ASV_25 | *Proteobacteria_Alphaproteobacteria_Rhodobacterales_Rhodobacteraceae_Cereibacter* | 0.01 | 0.00 | 3.33 | 0.10 | 0.18 | 0.00 | 0.73 | 1.40 | 0.80 |
| ASV_26 | *Cyanobacteria_Melainabacteria_Obscuribacterales* | 0.00 | 0.01 | 0.01 | 0.01 | 0.99 | 0.17 | 3.56 | 1.44 | 1.34 |
| ASV_29 | *Proteobacteria_Alphaproteobacteria_Caulobacterales_Hyphomonadaceae* | 0.00 | 0.00 | 0.06 | 0.01 | 6.21 | 0.92 | 3.12 | 0.72 | 0.70 |
| ASV_30 | *Planctomycetes_Planctomycetacia_Planctomycetales* | 0.00 | 0.00 | 0.23 | 0.00 | 0.22 | 0.01 | 2.00 | 0.40 | 0.28 |
| ASV_31 | *Proteobacteria_Gammaproteobacteria_Pseudomonadales_Moraxellaceae_Acinetobacter* | 0.00 | 0.00 | 0.00 | 0.00 | 0.13 | 0.02 | 0.22 | 0.17 | 0.01 |
| ASV_32 | *Proteobacteria_Alphaproteobacteria_Rhizobiales_Hyphomicrobiaceae_Hyphomicrobium* | 0.00 | 0.00 | 0.00 | 0.00 | 0.07 | 0.09 | 0.01 | 2.62 | 1.86 |
| ASV_33 | *Proteobacteria_Alphaproteobacteria_Sphingomonadales_Sphingomonadaceae_Sphingomonas* | 0.00 | 0.00 | 0.00 | 0.00 | 0.46 | 0.00 | 0.00 | 2.28 | 1.43 |
| ASV_34 | *Proteobacteria_Gammaproteobacteria_Betaproteobacteriales_Gallionellaceae* | 0.28 | 0.00 | 0.00 | 0.00 | 0.00 | 0.00 | 0.01 | 0.28 | 0.29 |
| ASV_36 | *Planctomycetes_Phycisphaerae_Phycisphaerales_Phycisphaeraceae* | 0.00 | 0.00 | 0.18 | 0.00 | 1.23 | 1.29 | 2.52 | 0.73 | 0.64 |
| ASV_37 | *Proteobacteria_Gammaproteobacteria_Betaproteobacteriales_Sulfuricellaceae_Sulfuricella* | 0.05 | 0.00 | 0.00 | 0.00 | 0.10 | 0.00 | 0.00 | 1.70 | 1.54 |
| ASV_40 | *Proteobacteria_Alphaproteobacteria_Rickettsiales* | 0.00 | 0.02 | 0.02 | 0.01 | 5.00 | 6.43 | 1.03 | 0.19 | 0.30 |
| ASV_41 | *Proteobacteria_Alphaproteobacteria_Rickettsiales* | 0.00 | 0.00 | 0.00 | 0.00 | 4.23 | 8.53 | 1.02 | 0.48 | 0.44 |
| ASV_44 | *Bacteroidetes_Flavobacteriia_Flavobacteriales_Flavobacteriaceae_Flavobacterium* | 0.22 | 0.87 | 0.26 | 0.75 | 0.02 | 0.00 | 0.81 | 0.01 | 0.00 |
| ASV_80 | *Proteobacteria_Alphaproteobacteria_Rhizobiales_Beijerinckiaceae_Methylobacterium* | 0.00 | 0.00 | 0.00 | 0.01 | 0.04 | 1.70 | 4.66 | 0.87 | 0.53 |

**Table S7**: Pair-wise beta diversity comparisons between corresponding locations from each system

|  | **Structure based metrics** | | | | **Membership based metrics** | | | |
| --- | --- | --- | --- | --- | --- | --- | --- | --- |
|  | **Bray-Curtis** | | **Weighted UniFrac** | | **Jaccard** | | **Unweighted UniFrac** | |
| **Sample comparison** | **Mean** | **SD** | **Mean** | **SD** | **Mean** | **SD** | **Mean** | **SD** |
| R_SW vs S_SW | 0.71 | 0.10 | 0.47 | 0.13 | 0.84 | 0.06 | 0.71 | 0.06 |
| R_FI vs S_FI | 0.49 | 0.11 | 0.31 | 0.09 | 0.66 | 0.10 | 0.56 | 0.10 |
| R_FB vs S_FB | 0.66 | 0.08 | 0.42 | 0.13 | 0.75 | 0.05 | 0.63 | 0.03 |
| R_FE vs S_FE | 0.48 | 0.13 | 0.34 | 0.16 | 0.58 | 0.08 | 0.50 | 0.04 |
| R_CHLA vs S_CHLA | 0.72 | 0.20 | 0.60 | 0.24 | 0.92 | 0.04 | 0.82 | 0.07 |
| R_CHLB vs S_CHLB | 0.72 | 0.24 | 0.51 | 0.32 | 0.93 | 0.06 | 0.82 | 0.10 |
| R_CHM vs S_CHM | 0.74 | 0.21 | 0.67 | 0.16 | 0.82 | 0.06 | 0.69 | 0.05 |
| R_DS1 vs S_DS1 | 0.72 | 0.23 | 0.67 | 0.11 | 0.83 | 0.08 | 0.72 | 0.04 |
| R_DS2 vs S_DS2 | 0.67 | 0.17 | 0.56 | 0.10 | 0.83 | 0.06 | 0.72 | 0.04 |

**Table S8.** Permutation test for distance-based redundancy analysis (dbRDA) under reduced model

|  | SumOfSqs | F | Pr(>F) |
| --- | --- | --- | --- |
| Conductivity | 0.382 | 1.1569 | 0.2335 |
| DOC | 0.58 | 1.7565 | 0.0233* |
| Free Cl2 | 0.762 | 2.3074 | 0.0021** |
| Total Cl2 | 0.241 | 0.73 | 0.8524 |
| Monochloramine | 0.645 | 1.954 | 0.0095** |
| Alkalinity | 0.515 | 1.559 | 0.0461* |
| Ammonium | 0.972 | 2.9447 | 0.0001*** |
| Nitrite | 0.391 | 1.1849 | 0.2105 |
| Nitrate | 0.379 | 1.1487 | 0.2421 |
| pH | 0.37 | 1.121 | 0.2779 |
| Temperature | 0.697 | 2.1121 | 0.0062** |
| Turbidity | 0.207 | 0.6271 | 0.9505 |

Signif. codes: 0 ‘***’ 0.001 ‘**’ 0.01 ‘*’ 0.05 ‘.’ 0.1 ‘ ’ 1

**Table S9.** Variance Partition analyses using water chemistry/environmental parameters identified as significant being significantly associated with Bray-Curtis distances by dbRDA analyses.

Explanatory tables:

X1 = Ammonium, X2 = Free_Cl2, X3 = Temp, X4 = Monochloramine

No. of explanatory tables: 4

Total variation (SS): 40.502

No. of observations: 111

| **Variables and their combinations** | **Degrees of Freedom** | **R square** | **Adjusted R square** | **Testable** |
| --- | --- | --- | --- | --- |
| [aeghklno] = X1 | 1 | 0.04908 | 0.04036 | TRUE |
| [befiklmo] = X2 | 1 | 0.05648 | 0.04782 | TRUE |
| [cfgjlmno] = X3 | 1 | 0.02175 | 0.01277 | TRUE |
| [dhijkmno] = X4 | 1 | 0.04074 | 0.03194 | TRUE |
| [abefghiklmno] = X1+X2 | 2 | 0.07773 | 0.06065 | TRUE |
| [acefghjklmno] = X1+X3 | 2 | 0.07754 | 0.06046 | TRUE |
| [adeghijklmno] = X1+X4 | 2 | 0.07341 | 0.05625 | TRUE |
| [bcefgijklmno] = X2+X3 | 2 | 0.07574 | 0.05862 | TRUE |
| [bdefhijklmno] = X2+X4 | 2 | 0.07748 | 0.0604 | TRUE |
| [cdfghijklmno] = X3+X4 | 2 | 0.06379 | 0.04646 | TRUE |
| [abcefghijklmno] = X1+X2+X3 | 3 | 0.10414 | 0.07902 | TRUE |
| [abdefghijklmno] = X1+X2+X4 | 3 | 0.10786 | 0.08284 | TRUE |
| [acdefghijklmno] = X1+X3+X4 | 3 | 0.10194 | 0.07676 | TRUE |
| [bcdefghijklmno] = X2+X3+X4 | 3 | 0.09554 | 0.07018 | TRUE |
| [abcdefghijklmno] = All | 4 | 0.12838 | 0.09549 | TRUE |
| **Individual fractions** | |  |  |  |
| [a] = X1 \| X2+X3+X4 | 1 |  | 0.02531 | TRUE |
| [b] = X2 \| X1+X3+X4 | 1 |  | 0.01873 | TRUE |
| [c] = X3 \| X1+X2+X4 | 1 |  | 0.01265 | TRUE |
| [d] = X4 \| X1+X2+X3 | 1 |  | 0.01647 | TRUE |
| [e] | 0 |  | 0.00499 | FALSE |
| [f] | 0 |  | 0.00786 | FALSE |
| [g] | 0 |  | -0.00287 | FALSE |
| [h] | 0 |  | -0.00491 | FALSE |
| [i] | 0 |  | -0.00017 | FALSE |
| [j] | 0 |  | 0.00572 | FALSE |
| [k] | 0 |  | 0.0223 | FALSE |
| [l] | 0 |  | -0.00312 | FALSE |
| [m] | 0 |  | -0.00613 | FALSE |
| [n] | 0 |  | -0.0047 | FALSE |
| [o] | 0 |  | 0.00336 | FALSE |
| [p] = Residuals | 0 |  | 0.90451 | FALSE |
| **Controlling 2 tables X** | |  |  |  |
| [ae] = X1 \| X3+X4 | 1 |  | 0.0303 | TRUE |
| [ag] = X1 \| X2+X4 | 1 |  | 0.02244 | TRUE |
| [ah] = X1 \| X2+X3 | 1 |  | 0.0204 | TRUE |
| [be] = X2 \| X3+X4 | 1 |  | 0.02372 | TRUE |
| [bf] = X2 \| X1+X4 | 1 |  | 0.02659 | TRUE |
| [bi] = X2 \| X1+X3 | 1 |  | 0.01856 | TRUE |
| [cf] = X3 \| X1+X4 | 1 |  | 0.02051 | TRUE |
| [cg] = X3 \| X2+X4 | 1 |  | 0.00978 | TRUE |
| [cj] = X3 \| X1+X2 | 1 |  | 0.01837 | TRUE |
| [dh] = X4 \| X2+X3 | 1 |  | 0.01156 | TRUE |
| [di] = X4 \| X1+X3 | 1 |  | 0.0163 | TRUE |
| [dj] = X4 \| X1+X2 | 1 |  | 0.02219 | TRUE |
| **Controlling 1 table X** | |  |  |  |
| [aghn] = X1 \| X2 | 1 |  | 0.01283 | TRUE |
| [aehk] = X1 \| X3 | 1 |  | 0.04768 | TRUE |
| [aegl] = X1 \| X4 | 1 |  | 0.02431 | TRUE |
| [bfim] = X2 \| X1 | 1 |  | 0.02029 | TRUE |
| [beik] = X2 \| X3 | 1 |  | 0.04585 | TRUE |
| [befl] = X2 \| X4 | 1 |  | 0.02846 | TRUE |
| [cfjm] = X3 \| X1 | 1 |  | 0.02009 | TRUE |
| [cgjn] = X3 \| X2 | 1 |  | 0.0108 | TRUE |
| [cfgl] = X3 \| X4 | 1 |  | 0.01452 | TRUE |
| [dijm] = X4 \| X1 | 1 |  | 0.01589 | TRUE |
| [dhjn] = X4 \| X2 | 1 |  | 0.01258 | TRUE |
| [dhik] = X4 \| X3 | 1 |  | 0.03368 | TRUE |

**Supplementary figures**

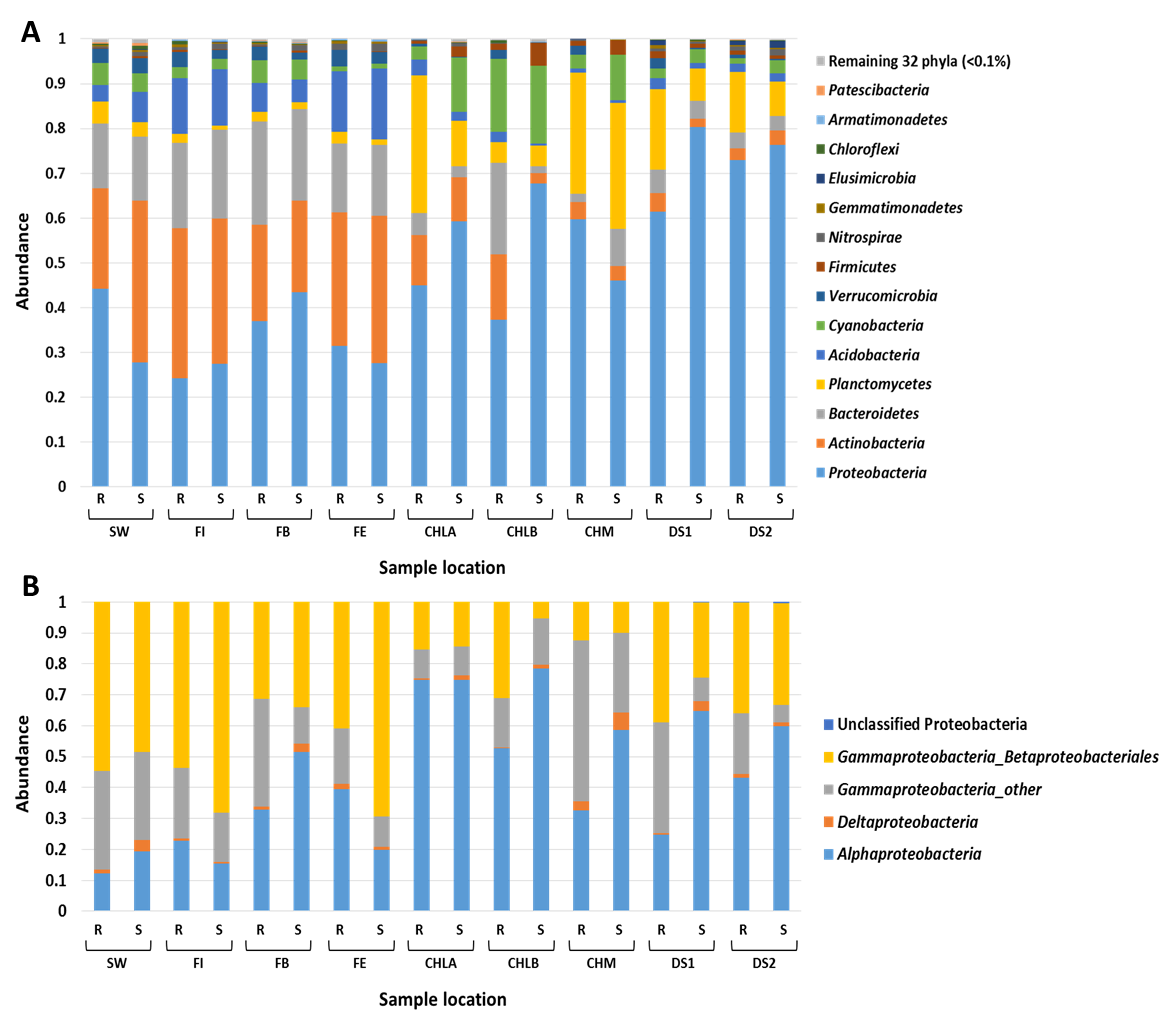

**Fig. S1**: (A) Phylum-level mean relative abundance of bacterial sequences detected over the duration of the study at each sample location within the two DWTPs and corresponding DWDS (R and S DWDS sections). The 14 most abundant and unclassified phyla (> 0.1%) are shown here, with the remaining 32 phyla (< 0.1%) grouped together as a single group. Phyla are shown in the legend on the right of the figure. See Table S3 for mean relative abundances. (B) Mean relative abundance of proteobacterial classes detected over the duration of the study at each sample location for each system.

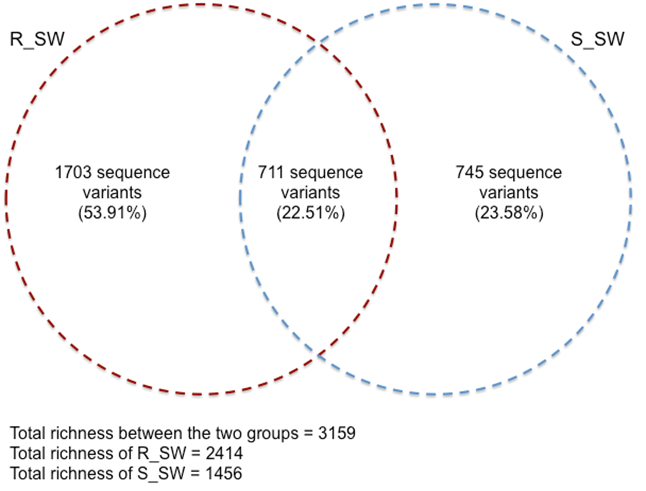

**Fig. S2**: Venn diagram showing the shared amplicon sequence variants (ASVs) between the two source waters.

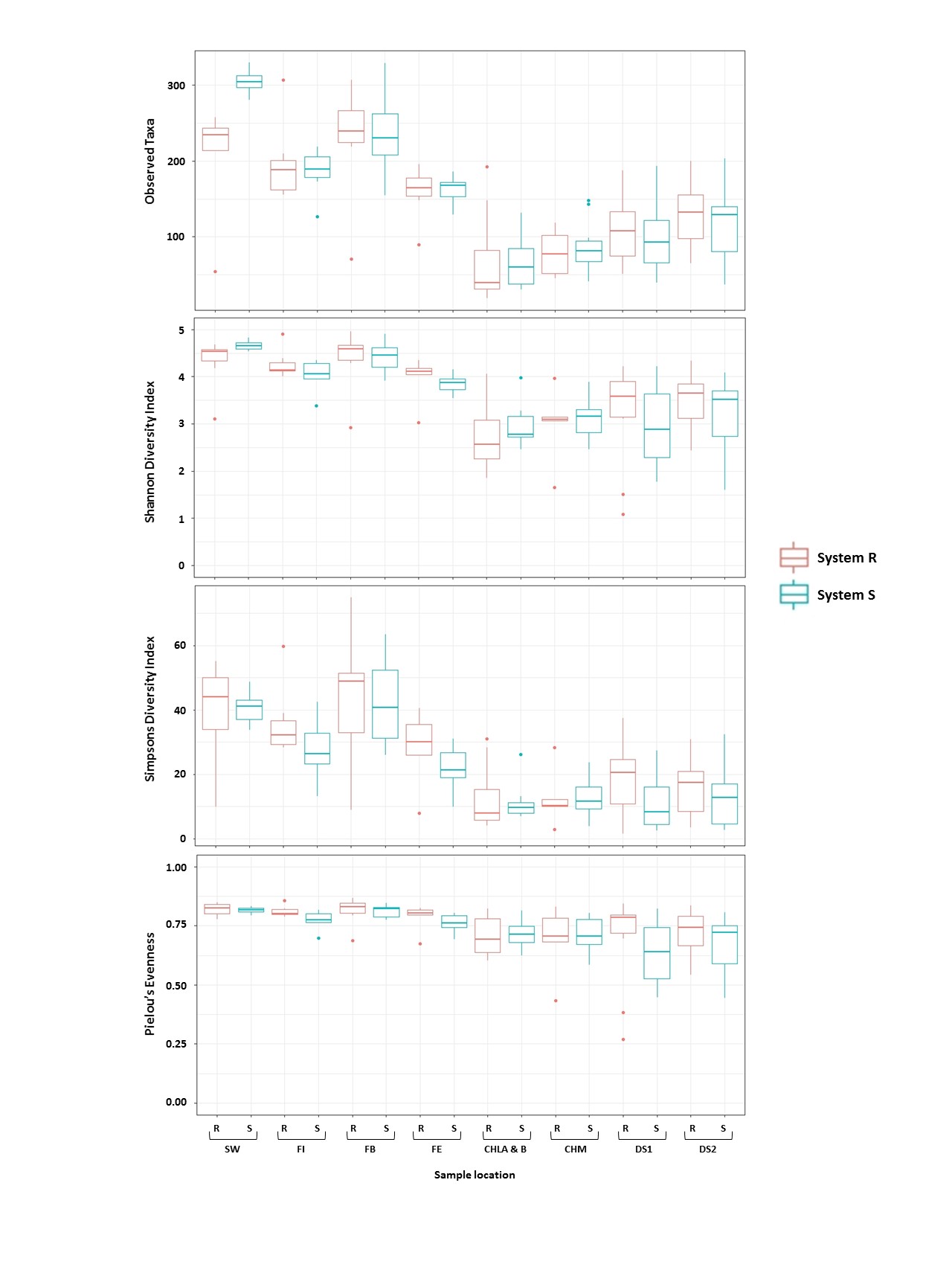
**Fig. S3**: Spatial changes in richness (observed taxa), diversity (Shannon Diversity Index and Inverse Simpson Diversity Index) and evenness (Pielou’s evenness) averaged across all sampling locations for each month. Points represent all sample sites collected for each month. Samples colored based on DWTP and corresponding DWDS (Lines R and S) (subsampled at 1263 iterations=1000).

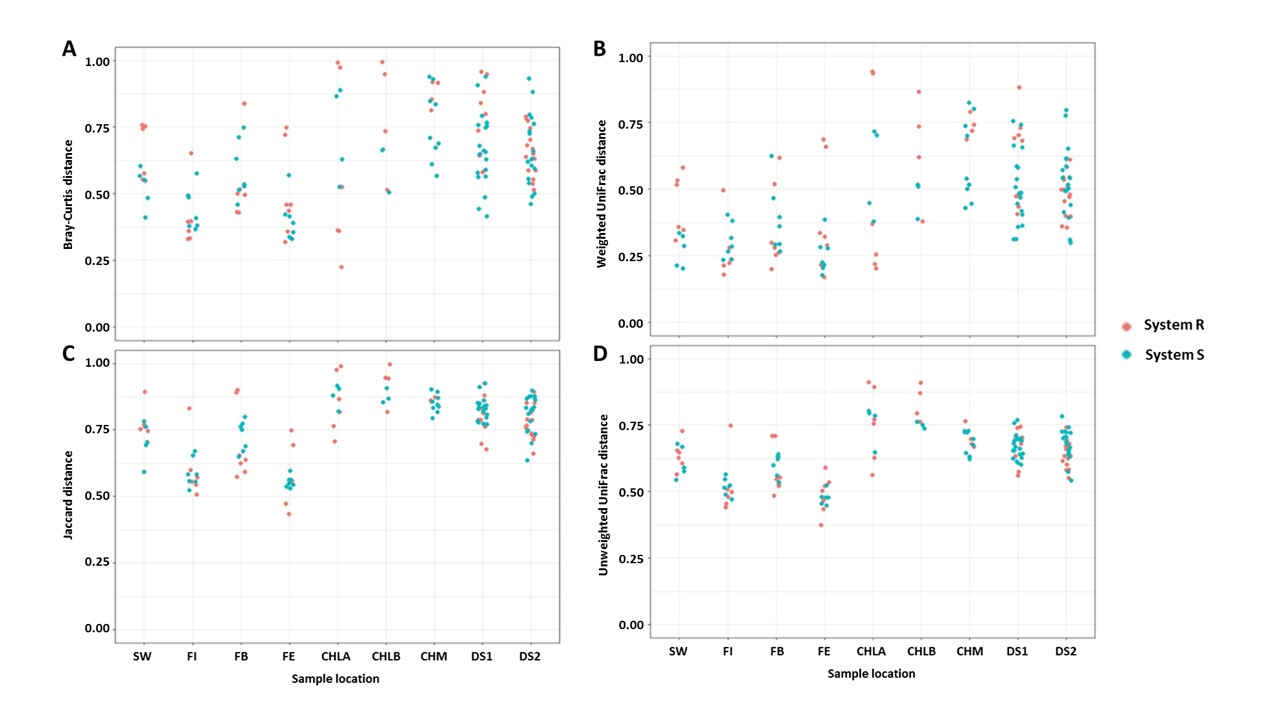

**Fig. S4** Temporal variation within each sample location from each system. Beta diversity pair-wise comparisons include samples from consecutive months within each location over the eight month study period for both structure-based metrics: (A) Bray-Curtis, (B) Weighted UniFrac and membership-based metrics: (C) Jaccard, (D) Unweighted UniFrac. Samples form System R are indicated in red and samples from System S are indicated in blue.

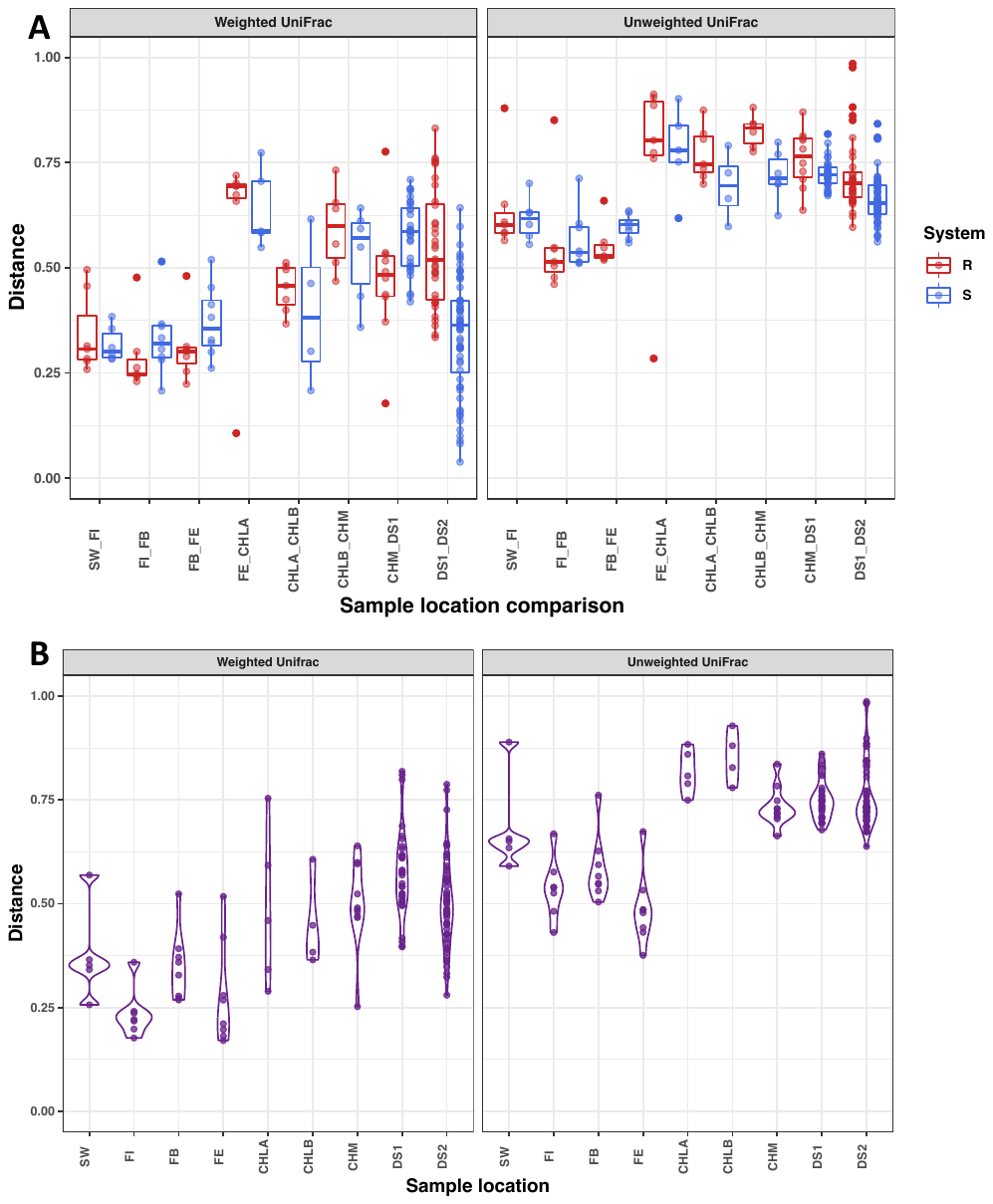

**Fig S5**: (A) Average pairwise beta diversity comparisons of both weighted and unweighted UniFrac analyses. Comparisons are between consecutive locations within each of the two systems for corresponding months. Sample abbreviations on the x-axis refer to comparisons of samples following the flow of bulk water through both systems, i.e. source water and filter inflow (SW_FI), filter inflow and filter bed media (FI_FB), filter bed media and filter effluent (FB_FE), filter effluent and chlorinated water leaving the DWTP (FE_CHLA), chlorinated water leaving the DWTP and chlorinated water entering the secondary disinfection boosting station (CHLA_CHLB), chlorinated water entering the secondary disinfection boosting station and chloraminated water (CHLB_CHM), chloraminated water and distribution system site 1 (CHM_DS1) and finally distribution system site 1 and distribution system site 2 (DS1_DS2). Sample comparisons from System R indicated in red and System S in blue. (B) Direct pairwise beta diversity comparisons (weighted and unweighted UniFrac) between corresponding sampling locations from the two systems. Pairwise beta diversity comparisons include samples from the same month. Sample abbreviations on the x-axis refer to comparisons between the source waters (SW), filter inflows (FI), filter bed medias (FB), filter effluents (FE), chlorinated waters leaving the DWTP (CHLA), chlorinated waters entering the secondary disinfection boosting station (CHLB), chloraminated waters (CHM), distribution system sites 1 (DS1) and distribution system sites 2 (DS2). Mean and standard deviations of each comparison is shown in Table S7.

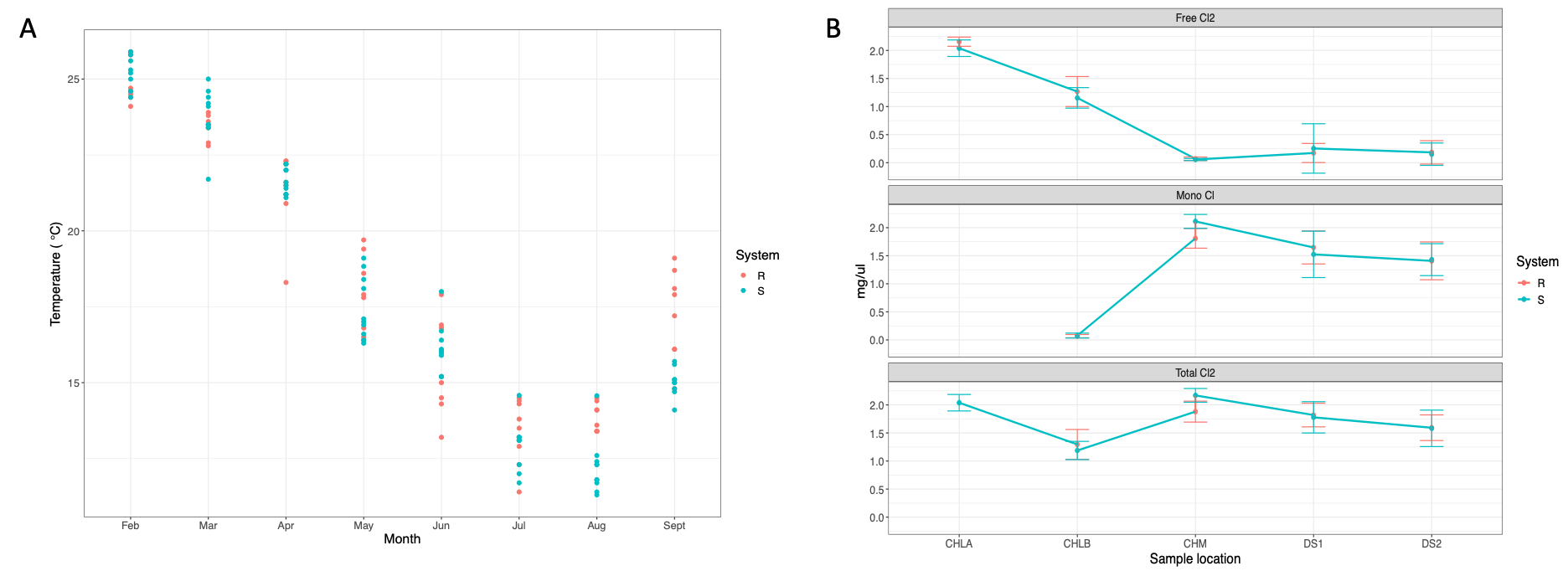

**Fig. S6.** Variation in temperature over the course of the study (8 months).

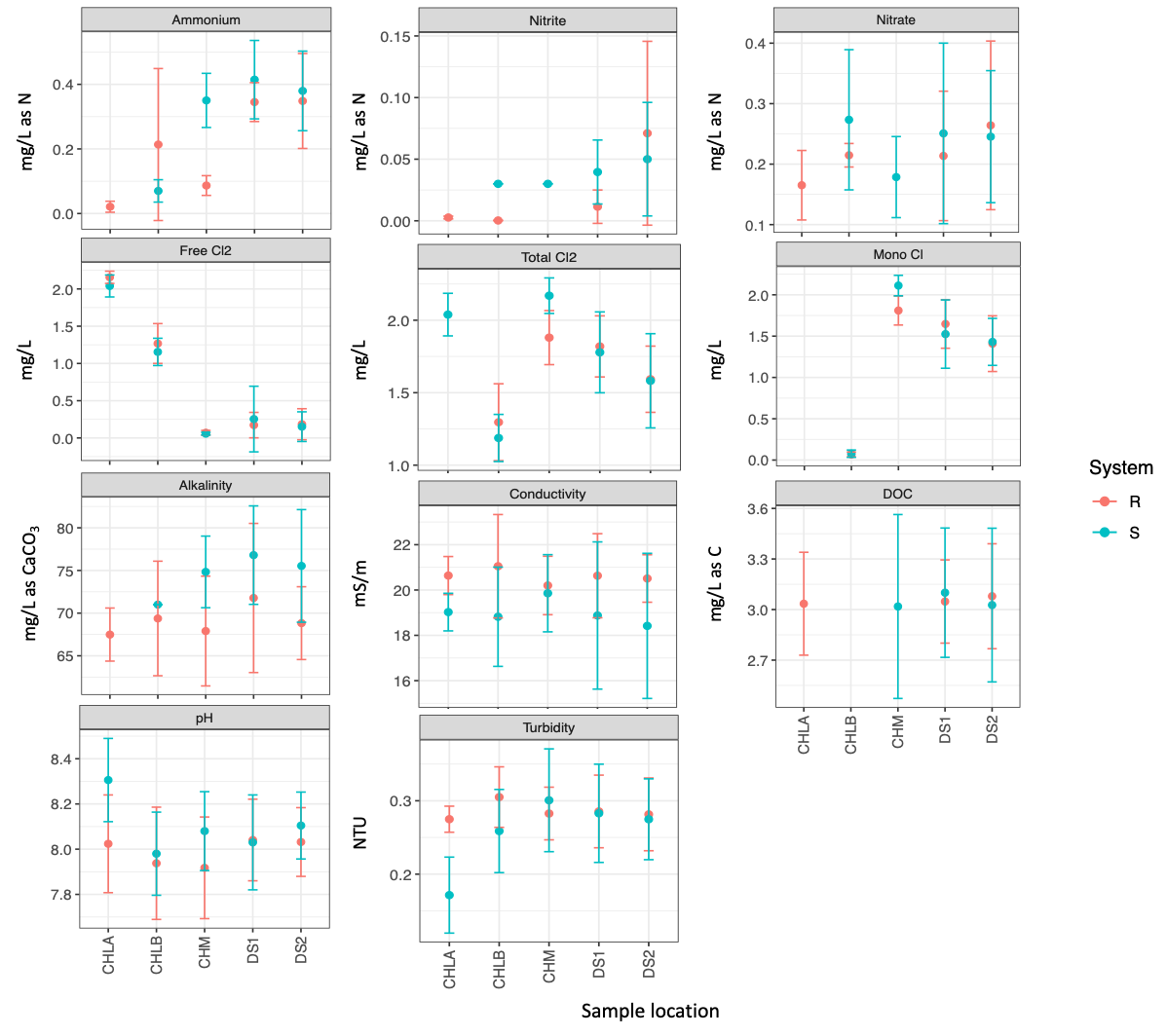

**Fig S7.** Variation in water quality parameters measured throughout the duration of the study for each location for both systems.

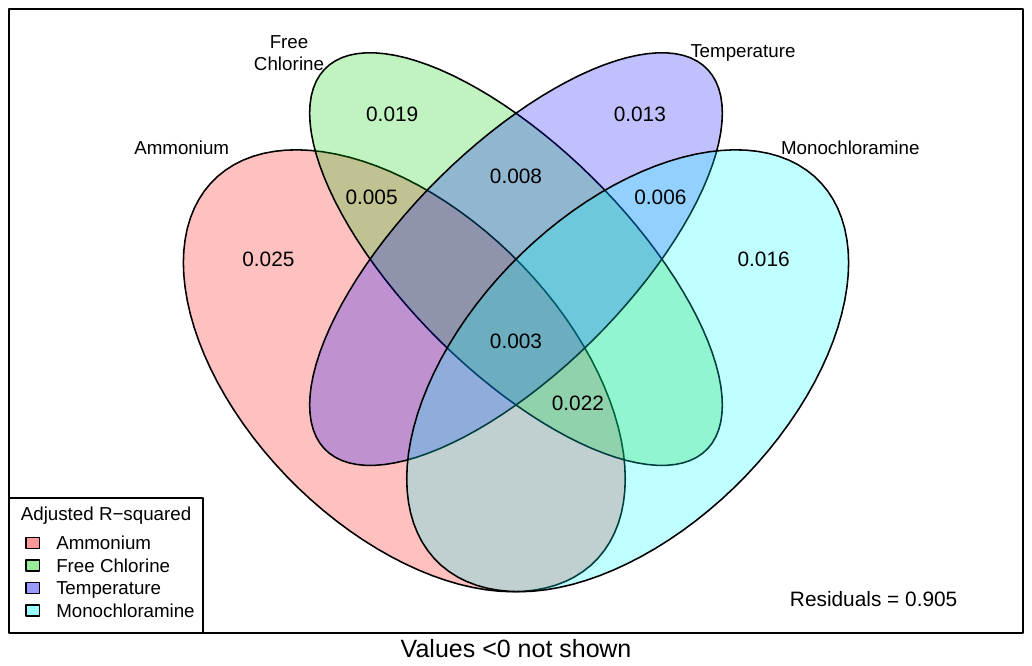

**Fig. S8. V**enn diagram indicates the faction of variation explained by the environmental parameters identified as significantly associated with ASV count-based Bray-Curtis distance matrices (Varpart analysis).
